# De novo gene synthesis by an antiviral reverse transcriptase

**DOI:** 10.1101/2024.05.08.593200

**Authors:** Stephen Tang, Valentin Conte, Dennis J. Zhang, Rimantė Žedaveinytė, George D. Lampe, Tanner Wiegand, Lauren C. Tang, Megan Wang, Matt W.G. Walker, Jerrin Thomas George, Luke E. Berchowitz, Marko Jovanovic, Samuel H. Sternberg

**Author notes:** These authors contributed equally to this work (Valentin Conte, Dennis J. Zhang). Present address: Department of Microbiology and Cell Biology, Montana State University, Bozeman, MT, USA.

## Abstract

Bacteria defend themselves from viral infection using diverse immune systems, many of which sense and target foreign nucleic acids. Defense-associated reverse transcriptase (DRT) systems provide an intriguing counterpoint to this immune strategy by instead leveraging DNA synthesis, but the identities and functions of their DNA products remain largely unknown. Here we show that DRT2 systems execute an unprecedented immunity mechanism that involves de novo gene synthesis via rolling-circle reverse transcription of a non-coding RNA (ncRNA). Unbiased profiling of RT-associated RNA and DNA ligands in DRT2-expressing cells revealed that reverse transcription generates concatenated cDNA repeats through programmed template jumping on the ncRNA. The presence of phage then triggers second-strand cDNA synthesis, leading to the production of long double-stranded DNA. Remarkably, this DNA product is efficiently transcribed, generating messenger RNAs that encode a stop codon-less, never-ending ORF (*neo*) whose translation causes potent growth arrest. Phylogenetic analyses and screening of diverse DRT2 homologs further revealed broad conservation of rolling-circle reverse transcription and Neo protein function. Our work highlights an elegant expansion of genome coding potential through RNA-templated gene creation, and challenges conventional paradigms of genetic information encoded along the one-dimensional axis of genomic DNA.

**One-Sentence Summary:** Bacterial reverse transcriptases synthesize extrachromosomal genes via rolling-circle amplification to confer potent antiviral immunity.

## INTRODUCTION

Mobile genetic elements (MGEs), including viruses, plasmids, and transposons, act as a major driving force to shape genome evolution by expressing protein enzymes that catalyze diverse DNA rearrangements (*1*). Many of these enzymes are encoded by the most abundant gene family in nature (*2*), and over sufficiently long timescales, MGEs can become dominant constituents of their host genomes due to their proliferative properties (*3*). In response, cells harbor arsenals of defense mechanisms to counter the pervasive spread of MGEs, which range in complexity from single-gene modules to the multi-organ vertebrate immune system (*4*). Strikingly, MGEs themselves often provide the source material from which anti-MGE mechanisms are derived (*5*). In vertebrates, for example, domestication of an ancestral transposon led to the evolution of V(D)J recombination, laying the groundwork for protein-based adaptive antiviral immunity by facilitating antigen receptor diversification (*6*). And in bacteria, recurrent exaptation of transposon-encoded genes enabled the emergence of CRISPR–Cas systems, which exploit guide RNA molecules to recognize and cleave foreign targets during nucleic acid-based adaptive antiviral immunity (*7–9*). MGEs have thus contributed to the perennial host–parasite arms race as both aggressor and defender, across all domains of life.

Inspired by the innovative molecular mechanisms of immunity born out of MGE co-option, we set out to investigate additional host pathways used by bacteria to defend against genetic invaders, and in particular, bacteriophages (*10*, *11*). We were especially intrigued by recent discoveries of diverse antiphage defense systems that encode reverse transcriptase (RT) enzymes (*12*, *13*). In stark contrast to well-studied phage defense pathways that target and destroy foreign nucleic acids using nucleases, such as restriction-modification (RM) and CRISPR–Cas, these RT-based systems presumably confer immunity via nucleic acid synthesis.

Prokaryotic RTs are thought to all descend from a common retroelement ancestor, the Group II intron, which catalyzes self-splicing and site-specific DNA insertion, and is also thought to be the precursor to the eukaryotic spliceosome (*14*). The antiviral roles of multiple classes of domesticated retroelements have emerged in recent years (*14*), reinforcing the notion that genetic conflicts position MGEs in both offensive and defensive roles. One class, for example, comprises fusions or operonic associations between an RT and the Cas1 integrase, itself a domesticated transposase, to record molecular memories from past infections by writing fragments of viral RNA into the CRISPR DNA (*15–17*). Retrons constitute another class that mediate innate antiphage immunity using complex operons that typically consist of an RT domain, a non-coding RNA (ncRNA), and a toxin protein (*12*, *13*, *18*). In uninfected cells, the RT reverse transcribes the ncRNA to form a complementary DNA (cDNA) product, which is thought to maintain the toxin in an inactive state; phage infection then drives cDNA modification, leading to activation of the toxin and initiation of cell suicide (*12*, *18*). This general strategy, known as abortive infection, prevents the invading phage from completing its lytic cycle and provides population-level immunity at the expense of the infected cell.

Defense-associated RT (DRT) systems comprise a third class of retroelements with antiphage function, which originate from a monophyletic clade of RTs termed the Unknown Group (UG) (*13*, *19*). In contrast to RT-Cas1 and retron systems, many DRTs feature single-gene operons (*19*), implying that reverse transcription activity alone could be responsible for providing phage defense. Initial experimental efforts have confirmed the phage defense functions of 9 UG subgroups (DRT1-9), in addition to demonstrating the functional requirement for an intact RT domain (*13*, *19*). However, the identities of the DRT cDNA products, and their mechanisms of immunity, have not been studied.

Here we develop a systematic approach to profile the cDNA products of any RT enzyme of interest, and apply it to DRT2-family phage defense systems. This strategy revealed an unprecedented rolling-cir-cle reverse transcription activity, leading to the production of repeated, concatenated cDNA products. Remarkably, we discover that these cDNAs contain an open reading frame (ORF) that remains in-frame through each repeat, and that promoter sequences formed across the repeat junction result in abundant mRNA transcription. Expression of the never-ending ORF (*neo*) gene, which is stringently regulated by the presence or absence of phage, leads to rapid growth arrest and programmed dormancy. Beyond revealing an elegant example of retroelement exaptation for host–MGE defense, we present evidence supporting the broad conservation of this mechanism for RNA-templated creation of extrachromosomal genes.

## RESULTS

### cDNA synthesis by a defense-associated reverse transcriptase

We focused our attention on DRT2 systems because of their intriguing minimal architecture, consisting of a single ORF and upstream ncRNA described previously (*19*). Unlike most other DRT and retron systems, which typically encode a reverse transcriptase enzyme alongside one or more additional protein domains predicted to function as effectors of the immune response, DRT2 systems lack additional protein-coding genes, and the RT protein lacks domains beyond the predicted RNA-directed DNA polymerase (*11*, *19*) (**fig. S1A**). We therefore hypothesized that the cDNA product of the RT enzyme likely plays a critical and central role in the DRT2 immune mechanism. To identify this cDNA, we developed a sequencing approach to systematically identify RT-associated cDNA synthesis products based on immunoprecipitation of FLAG-RT fusions, which we termed cDNA immunoprecipitation and sequencing (cDIP-seq) (**Materials and Methods**), and performed these experiments alongside traditional RNA immunoprecipitation (RIP)-seq to also capture RT-associated RNA substrates (**Fig. 1A and fig. S1B**). We validated this approach on the well-studied Retron-Eco1 (formerly Ec86) (*20*) after first verifying that the FLAG-tagged retron RT retained phage defense activity comparable to the WT RT (**fig. S1C**). RIPseq and cDIP-seq of Retron-Eco1 recapitulated all known features of both the msRNA and msDNA, including RNase H processing of msRNA, and precise 5′ and 3′ ends of the msDNA (*21*) (**fig. S1D,E**). These results increased our confidence that a similar approach could provide new insights into the DRT2 molecular mechanism, and so we turned our attention to a candidate system from *Klebsiella pneumoniae* (*Kpn*DRT2).

**Fig. 1.**
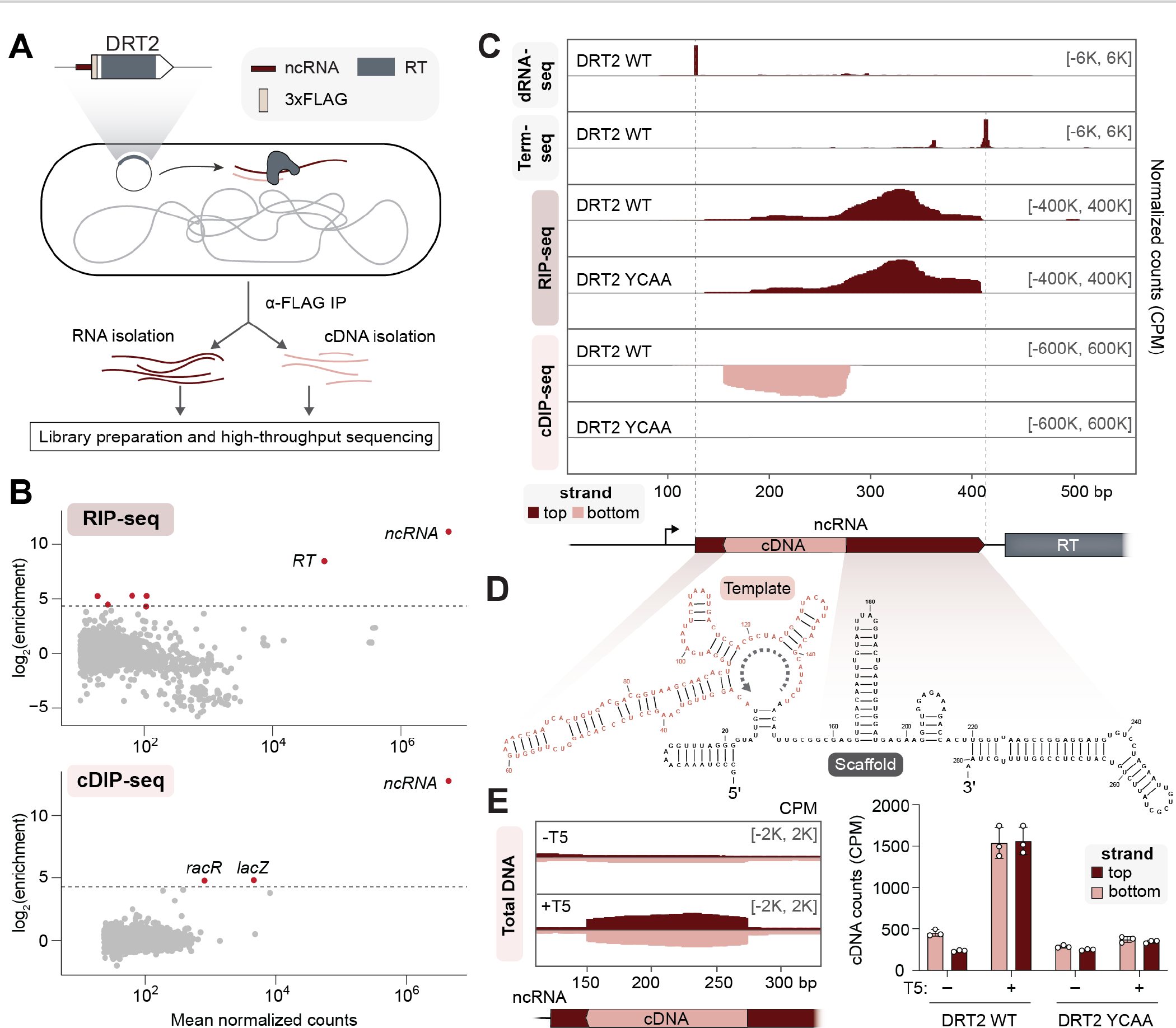
Systematic discovery of DRT2 reverse transcription substrates and products *in vivo*. (A) Schematic of RNA immunoprecipitation (RIP) and cDNA immunoprecipitation (cDIP) sequencing approaches to identify nucleic acid substrates of FLAG-tagged reverse transcriptase (RT) from *Kpn*DRT2. The plasmid-encoded immune system is schematized top left. **(B)** MA plots showing the RT-mediated enrichment of RNA (top) and DNA (bottom) loci from RIP-seq and cDIP-seq experiments, relative to input controls. Each dot represents a transcript, and red dots denote transcripts with > 20-fold enrichment and false discovery rate (FDR) < 0.05. **(C)** dRNA-seq, Term-seq, RIP-seq, and cDIP-seq coverage tracks, from top to bottom, for either WT RT or a catalytically inactive RT mutant (YCAA). dRNA-seq and Term-seq enrich RNA 5′ and 3′ ends, respectively, whereas RIP-seq and cDIP-seq identify RT-associated RNA and DNA ligands. Red and pink denote top and bottom strands, respectively, and the DRT2 locus is shown at bottom; coordinates are numbered from the beginning of the *K. pneumoniae*-derived sequence on the expression plasmid. Data are normalized for sequencing depth and plotted as counts per million reads (CPM). **(D)** Predicted secondary structure of the *Kpn*DRT2 ncRNA. The cDNA template region is colored in pink, and the gray dotted line denotes the direction of reverse transcription. **(E)** Coverage over the DRT2 ncRNA locus from total DNA sequencing of cells +/T5 phage infection (left), and bar graph of cDNA counts for the same samples alongside the YCAA mutant (right). Red and pink denote top and bottom strands, respectively; data are mean ± s.d. (n = 3).

After confirming that fusing the *Kpn*DRT2 RT with a FLAG epitope tag did not affect defense activity against T5 phage (**fig. S1C**), we performed RIP-seq and cDIP-seq from cells constitutively expressing plasmid-encoded ncRNA and RT from their native promoter, and then performed genome-wide analyses to identify RNA and cDNA molecules enriched by IP. The resulting datasets revealed that the highest enriched RNA and cDNA transcripts mapped to the *Kpn*DRT2 ncRNA locus (**Fig. 1B**), suggesting that the primary substrate for reverse transcription by DRT2 is encoded in *cis*, similar to retron systems. We ascribed the apparent RIP-seq enrichment of *RT* mRNA to the likely presence of read-through transcripts extending from the ncRNA into the coding sequence. Other enriched hits from cDIP-seq data were also found in control experiments using a catalytically inactive RT mutant (*13*) (hereafter YCAA), suggesting a spurious origin (**fig. S2A**). Interestingly, RIP-seq and cDIP-seq experiments in the presence of T5 phage also revealed a strong and specific enrichment of transcripts derived from the DRT2 ncRNA locus (**fig. S2B**), indicating that RT substrate choice is largely unchanged during phage infection.

Mapping of RIP-seq and cDIP-seq data onto the *Kpn*DRT2 locus revealed the presence of a large ncRNA and a seemingly well-defined cDNA with the opposite strandedness relative to the ncRNA, as expected for reverse transcription (**Fig. 1C**). Control experiments with the inactive YCAA RT mutant demonstrated that ncRNA enrichment occurred independently of reverse transcriptase activity, whereas cDNA enrichment from this locus required an intact RT active site (**Fig. 1C and fig. S2A**). We next leveraged custom RNA-seq library preparation protocols based on dRNA-seq (*22*) and Term-seq (*23*) to demarcate the precise 5′ and 3′ ends of the 281-nucleotide (nt) ncRNA (**Fig. 1C**), while analyzing start and end coordinates from cDIP-seq alignments to define the 5′ and 3′ ends of the 119-nt cDNA (**fig. S2C**). Interestingly, dRNA-seq data revealed a single transcription start site (TSS) upstream of the ncRNA, but not the *RT* gene (**Fig. 1C**), suggesting that the ncRNA and *RT* share an upstream promoter, and are separated into mature transcripts via an unknown processing step. We next used a multiple sequence alignment (MSA) of DRT2 homologs to generate a covariance model of the ncRNA (**fig. S2D**), which was in excellent agreement with the *Kpn*DRT2 ncRNA secondary structure predicted by *in silico* RNA folding (*24*) (**Fig. 1D**). The ncRNA features numerous conserved stem-loop (SL) elements, a template region corresponding to the cDNA product abutted by a short basal stem, and a large 3′ region that we hypothesize serves as a scaffold for sequence and/or structure-guided recruitment of the RT.

We next investigated how cDNA synthesis is altered as cells are actively infected by, and defending against, T5 phage. cDIP-seq data largely recapitulated the same observations made in the absence of phage (**fig. S2B,E**), but we were wary of drawing conclusions about reverse transcription output based on an approach that would only quantify cDNAs still bound to the RT after immunoprecipitation. We therefore turned to total DNA sequencing using the input controls from cDIP-seq experiments, and our analyses revealed a strong induction in *Kpn*DRT2 cDNA levels upon phage infection (**Fig. 1E**). Surprisingly, while cDNA synthesis products in the absence of phage were predominantly single-stranded, with opposite strandedness to the ncRNA, the presence of phage induced higher levels of both the initial cDNA product and its reverse complement (**Fig. 1E**). Although these experiments do not reveal the identity of the polymerase necessary for second-strand synthesis, we hypothesize that the RT likely possesses both RNA-templated and DNA-templated DNA polymerase activity, similar to other well-studied bacterial reverse transcriptases (*25*), and that conversion of ssDNA to dsDNA may be a key step within the antiviral defense pathway.

### Rolling-circle reverse transcription generates concatenated cDNA products

We next sought to investigate the sequence requirements and potential antiviral function of cDNA synthesis. We began by mutating the sequence of individual SLs throughout the ncRNA in order to eliminate base-pairing, focusing on SL1 at the 5′ end, SL2 at the base of the template region, SL5 within the template region, and SL6 within the scaffold region (**Fig. 2A**). Mutations to all four regions led to a complete loss of phage defense activity, indicating possible defects in ncRNA binding, cDNA synthesis, or both (**Fig. 2B**). When we directly interrogated ncRNA binding and cDNA synthesis by the RT using RIP-seq and cDIP-seq, respectively, we found that SL1 and SL6 mutants led to either a partial or complete loss of cDNA synthesis, likely due to disruptions in the positioning of the RT on the ncRNA (**Fig. 2C**). The SL5 mutant exhibited strong ncRNA and cDNA enrichment, as did an additional mutant in which the region surrounding the cDNA synthesis start site was scrambled (**fig. S3A-C**), suggesting that defense activity depends on not only cDNA synthesis, but also on the sequence of the cDNA product itself. The phenotype of the SL2 mutant, however, was puzzling: the sequence of the template region was completely unchanged and cDNA production resembled the WT system, and yet phage defense was completely abolished (**Fig. 2B,C**). This apparent discrepancy indicated that, beyond production of cDNA with the appropriate ncRNA-specified sequence, additional features of the cDNA product underlie phage defense activity.

**Fig. 2.**
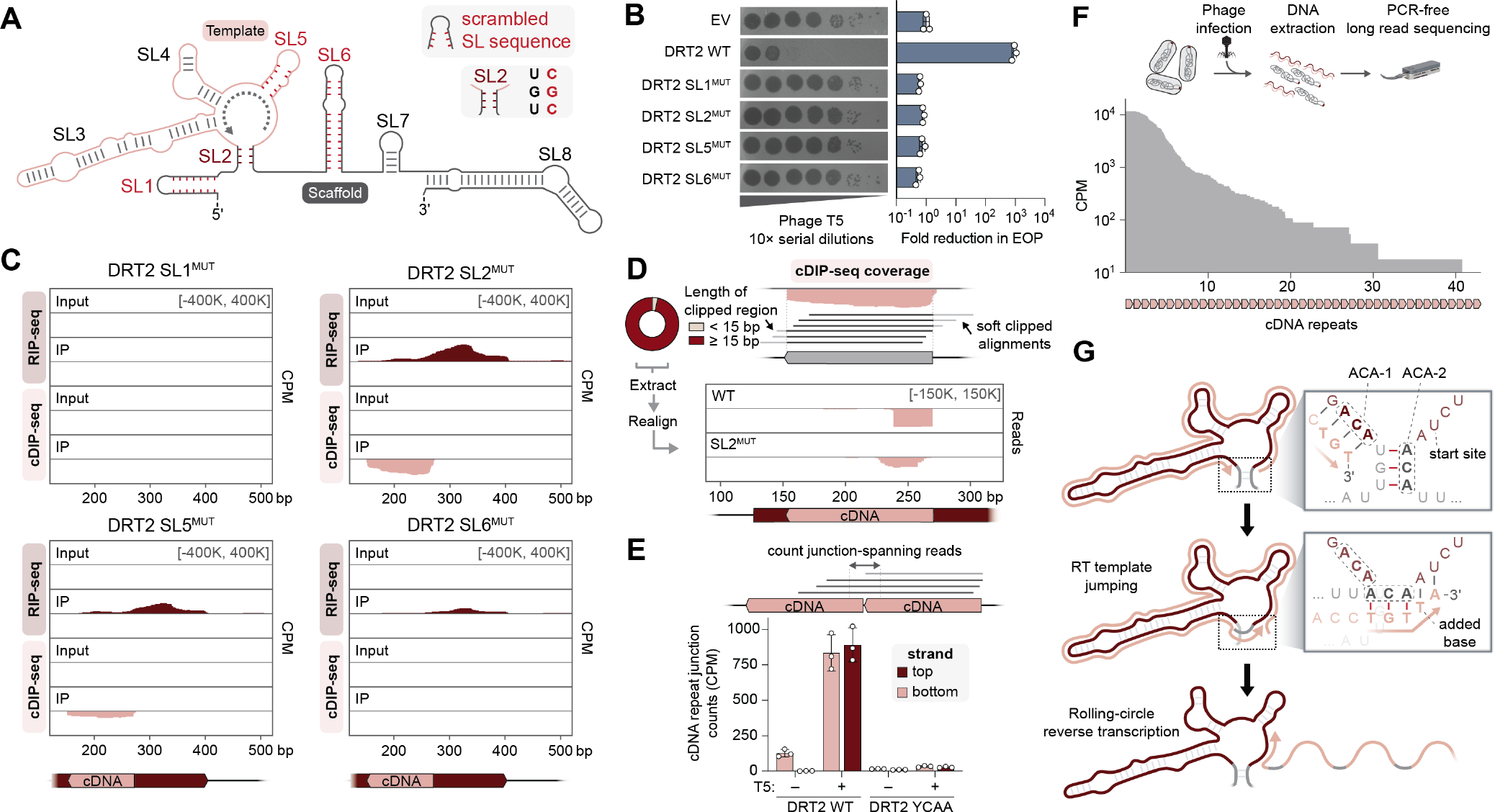
Rolling-circle reverse transcription generates a concatenated cDNA product. (A) Schematic of DRT2 ncRNA secondary structure, with stem-loops (SL) numbered 1–8 and selected perturbations highlighted in red. SL1^MUT^, SL5^MUT^, and SL6^MUT^ correspond to ncRNA mutants in which the SL bases were scrambled, resulting in the elimination of sequence motifs and secondary structure. SL2^MUT^ abolishes base pairing within the SL2 stem. Sequences of all mutants are presented in **Supplementary Table S3**. **(B)** Plaque assay showing loss of phage defense activity for all SL mutants from **A** (left), and bar graph quantifying the reduction in efficiency of plating (EOP, right); data are mean ± s.d. (n = 3). **(C)** RIP-seq and cDIP-seq coverage tracks for the indicated SL mutants alongside input controls, revealing a range of defects in either RNA binding, cDNA synthesis/binding, or both. **(D)** *Top*: Schematic of terminal portions of cDIP-seq reads (light gray) failing to align to the cDNA reference, resulting in ‘soft clipping’ and exclusion from coverage plots. A donut plot reporting the proportion of cDNA-mapping reads with the indicated lengths of 3′-clipped sequences is shown at left for DRT2 WT cDIP-seq. *Bottom*: Mapping of 3′-soft-clipped sequences from cDIP-seq experiments back to the DRT2 locus, demonstrating that they derive from the cDNA 5′ end. SL2^MUT^ exhibits an aberrant pattern relative to WT. **(E)** Schematic of sequencing reads that map across two concatenated cDNA repeats (top), and bar graph quantifying the abundance of junction-spanning reads from sequencing of total DNA in the indicated conditions (bottom). Red and pink denote top and bottom strands, respectively; data are mean ± s.d. (n = 3). **(F)** Schematic of long-read Nanopore sequencing workflow with DNA from phage-infected cells (top), and histogram of cDNA repeat length distribution for WT *Kpn*DRT2 from Nanopore sequencing (bottom). **(G)** Inferred mechanism of rolling-circle reverse transcription (RCRT) mediated by sequence and structural features of SL2. After synthesis of 5′-TGT-3′ templated by ACA-1 at the end of one cDNA repeat (top), the nascent DNA strand dissociates from its template and reanneals with the complementary ACA-2 following SL2 melting (middle). Template jumping initiates a subsequent round of reverse transcription, with concatenation of one cDNA repeat to the next and incorporation of one additional base at the repeat junction, ultimately leading to long rolling-circle cDNA products (bottom).

Given that the stem of SL2 borders the template region, we hypothesized that disruption of this structural element might lead to imprecise initiation or termination of cDNA synthesis, which in turn might explain the altered immune function. We inspected the 3′ termini of cDIP-seq reads more closely, and to our surprise, we found that the large majority of reads extended well beyond the boundary defined by the coverage signal; these extensions had been soft-clipped from the reads by conventional alignment algorithms (**Fig. 2D**). To determine the identity of these soft-clipped extensions, we extracted their sequences and mapped them back to the plasmid and *E. coli* genome. Remarkably, these sequences in fact derived from the 5′ end of the cDNA (**Fig. 2D**), suggesting a template jumping mechanism whereby the RT proceeds from the end of the template region back to the start, resulting in concatenated cDNA repeat products. Whereas the concatenated cDNAs generated by the WT system had a precise and uniform headto-tail junction, including one additional nucleotide immediately adjacent to SL2, junction sequences for the SL2 mutant were more heterogeneous (**Fig. 2D**). Indeed, when we quantified the frequency of the expected junction sequence across all tested ncRNA mutants, we found that only the SL5 and cDNA start mutants retained WT levels of the repeat junction, whereas all other SL mutants nearly eliminated the expected template jumping products (**fig. S3D**).

We next quantified concatenated cDNA products in total DNA samples from cells +/T5 phage infection. T5 phage infection triggered a large increase in the abundance of bottom-strand junction-spanning reads, corresponding to the initial products of RNA-templated DNA synthesis (**Fig. 2E**). This was matched with a concomitant increase in top-strand junction-spanning reads (**Fig. 2E**), suggesting that concatenated cDNA synthesis products are efficiently converted into dsDNA in a phage and RT-dependent manner. Similar analyses from cDIP-seq datasets showed a lesser decrease in top-strand junction-spanning reads during phage infection (**fig. S3E**), which we attribute to the RT likely having lower affinity for the dsDNA generated by second-strand cDNA synthesis, such that it releases these products in cells and/ or during immunoprecipitation.

Although short-read sequencing enabled accurate determination and quantification of cDNA repeat junctions, we next leveraged long-read Nanopore sequencing to assess the length of concatenated cDNA products, and to obtain orthogonal evidence of template jumping with a PCR-free approach. Remarkably, *Kpn*DRT2 cDNA products from phage-infected cells spanned a dramatic range of repeat lengths from 1–40 (**Fig. 2F**), revealing that reverse transcription by *Kpn*DRT2 is highly processive and involves many consecutive rounds of template jumping to generate long concatenated cDNA (from 120 to ∼5000 bp). Finally, we carefully inspected the sequence and secondary structure of the ncRNA in order to better understand the mechanism of template jumping. Concatenation of cDNA repeats occurs between the sequences directly abutting SL2, and we noticed that the terminal 3-nt of each repeat are templated by a conserved 3′- ACA-5′ (ACA-1) whose sequence perfectly matches the right half of SL2 (ACA-2; **Fig. 2G**). We therefore hypothesized that the RT may dynamically melt SL2 during each round of reverse transcription, allowing the terminal 5′-TGT-3′ of nascent cDNA transcripts to equilibrate between hybridization to ACA-1 and ACA-2, and thus prime a subsequent round of cDNA synthesis (**Fig. 2G**). This model was supported by a complete loss of defense activity in ncRNA mutants disrupting homology between the ACA motifs (**fig. S3F**). We note that the proposed cDNA concatenation mechanism resembles rolling-circle DNA replication (*26*), and henceforth refer to the generation of concatenated cDNA products as rolling-circle reverse transcription (RCRT; **Fig. 2G**).

### Concatenated cDNAs encode a translated open reading frame (ORF)

Conventional rolling-circle amplification utilizes a circular template, and thus we sought to rule out the alternative explanation that RCRT occurs as a result of ncRNA circularization at the repeat junction. To investigate this hypothesis, we reanalyzed our RIP-seq input controls, which represent total RNA-seq datasets, for the presence of reads spanning the repeat junction. Strikingly, we detected such reads abundantly in *Kpn*DRT2 samples, but they strictly depended on the presence of phage and an active RT, and even more unexpectedly, their strandedness was opposite to that of the ncRNA (**Fig. 3A**). This observation raised the intriguing possibility that cDNA second-strand synthesis might generate a template strand for another round of transcription by RNA polymerase (**Fig. 3B**). The resulting transcript would have opposite strandedness to the initial ncRNA and would contain multiple repeats of the cDNA sequence.

**Fig. 3.**
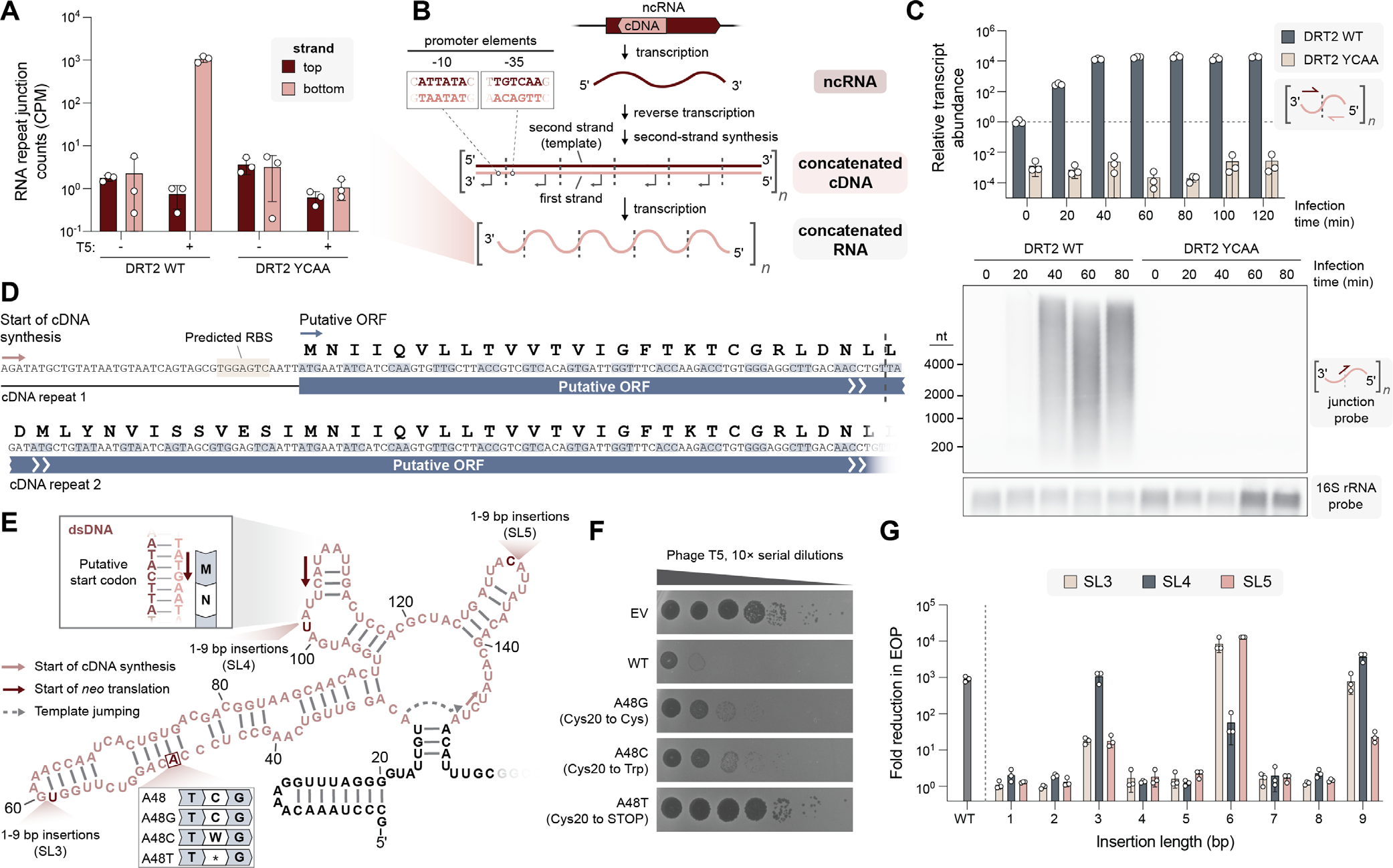
The concatenated cDNA product contains a never-ending ORF (*neo*). (A) Bar graph quantifying RNA-seq reads that map across two concatenated cDNA repeats, for the indicated conditions. Red and pink denote top and bottom strands, respectively; data are mean ± s.d. (n = 3). **(B)** Model showing the consecutive production of ncRNA (transcription), concatenated double-stranded cDNA (reverse transcription and second-strand synthesis), and concatenated RNA (transcription), all encoded by the DRT2 locus. Dashed lines indicate repeat–repeat junctions resulting from rolling-circle reverse transcription, and the inset (top left) shows the predicted promoter formed across the junction. **(C)** Bar graph quantifying relative concatenated RNA abundance in a phage infection time course experiment using RT-qPCR with repeat junction primers (top), and Northern blot of concatenated RNA using a junction-spanning probe (bottom). RT-qPCR data are normalized to WT uninfected cells (t = 0); data are mean ± s.d. (n = 3). **(D)** Putative open reading frame (ORF) encoded by the concatenated RNA. The start of cDNA synthesis and putative start of translation are indicated (pink and blue arrows, respectively), and the repeat–repeat junction is denoted with a dashed line. **(E)** Schematic of the cDNA template region (pink), with the putative start codon and experimentally tested mutations indicated. **(F)** Plaque assay showing that phage defense activity is eliminated with a single-bp substitution that introduces an in-frame stop codon, but is only modestly affected by synonymous or missense mutations. EV, empty vector. **(G)** Bar graph quantifying phage defense activity for insertions within SL3, SL4, or SL5, of the indicated length. Reduction in EOP is calculated relative to an EV control; data are mean ± s.d. (n = 3). The only mutants that retain phage defense activity have insertion lengths of a multiple of 3 bp.

In agreement with this hypothesis, inspection of the cDNA sequence produced by RCRT revealed consensus promoter elements spanning the repeat junction (**Fig. 3B**), highly reminiscent of transposon promoters that are selectively formed upon DNA circularization during the transposon excision step (*27*, *28*) These observations began to suggest that phage-induced second-strand cDNA synthesis might serve to trigger the production of a high-copy concatenated RNA molecule with downstream antiphage function.

Consistent with this idea, we found that concatenated RNAs were strongly induced shortly after phage infection by ∼10,000-fold, in a time-course infection experiment (**Fig. 3C**). Northern blot analysis from phage-infected cells using a probe selective for the chimeric junction revealed a broad size distribution spanning hundreds to thousands of nucleotides (**Fig. 3C**), in excellent agreement with the large size of cDNA products observed via Nanopore sequencing (**Fig. 2F**).

What could be the function of transcribing a repetitive cDNA sequence into RNA during an antiphage immune response? We closely examined the sequence of the cDNA and noticed that if we translated this sequence *in silico*, one out of three open reading frames (ORF) lacked any stop codons (**Fig. 3D and fig. S4A**). This observation led us to hypothesize that the concatenated RNA produced during phage infection might be translated to generate an antiviral polypeptide. This hypothesis was supported by the presence of a predicted ribosome binding site upstream of the predicted start codon (**Fig. 3D and fig. S4B**), and by the observation that programmed template jumping adds one additional nucleotide during each round of cDNA synthesis (**Fig. 2G**). This activity generates a 120-bp cDNA repeat unit comprising exactly 40 sense codons, such that the reading frame would be preserved through each repeat to yield a continuous ORF (**Fig. 3D**).

We set out to comprehensively test the hypothesis that translation of the continuous ORF within the concatenated RNA is necessary for phage defense. First, we identified a region within SL3 that was not strongly conserved in sequence, and introduced single-bp mutations that would generate a synonymous, missense, or nonsense codon (**Fig. 3E**). While the synonymous and missense mutations had mild effects on defense activity that we attributed to perturbation of the ncRNA secondary structure, the nonsense mutation completely abolished phage defense (**Fig. 3F**). We also mutated the predicted start codon and found that mutation to the non-canonical GUG start codon partially preserved defense activity, but all other mutations were inactive (**fig. S4C,D**). To assess whether translation of multiple contiguous repeats of the ORF was necessary for phage defense, we tested mutations that would introduce stop codons near the end of one full ORF repeat and found that these, too, led to a loss of defense (**fig. S4C,E**). Finally, inspired by classic experiments performed by Crick and Brenner to deduce the triplet nature of the genetic code (*29*), we selected three ncRNA loop regions and designed insertions ranging from 1–9 bp in length. We hypothesized that if translation of the ORF was required for defense, then out-of-frame mutations would lead to a loss of defense, where in-frame mutations (i.e., insertions of a multiple of 3 bp) would be partially or completely tolerated. Remarkably, we found that all out-of-frame perturbations across three non-conserved loop regions, including minimal 1-bp insertions, caused a >10^3^-fold decrease in phage defense, while insertions of 3, 6, or 9 bp maintained near-WT activity levels (**Fig. 3G**).

Collectively, these experiments provided compelling genetic evidence for the existence and expression of a cryptic gene produced by RNA-templated concatenation of DNA repeats. Intriguingly, this *de novo* gene exhibits a heterogeneous length distribution and lacks any in-frame stop codons, and thus we refer to it as *neo* (never-ending ORF).

### *Neo*-encoded polypeptides induce cell dormancy

Encouraged by our genetic assays supporting the translation of *neo*, we sought unambiguous biochemical evidence of Neo protein products. After analyzing the predicted amino acid composition of Neo, we designed a custom protease cocktail that would yield unique peptide fragments suitable for mass spectrometry (MS)-based proteomics (**fig. S5A**). We then extracted proteins from *Kpn*DRT2-expressing cells and performed liquid chromatography with tandem mass spectrometry (LC-MS/MS) analysis (**Fig. 4A**). Neo-derived peptides were exclusively detected in phage-infected cells that expressed the WT RT enzyme (**Fig. 4B**), and their abundances were substantial when compared to the rest of the *E. coli* proteome (**Fig. 4C**). These results provide concrete proof that *neo* mRNAs transcribed from concatenated cDNA genes are translated into protein.

**Fig. 4.**
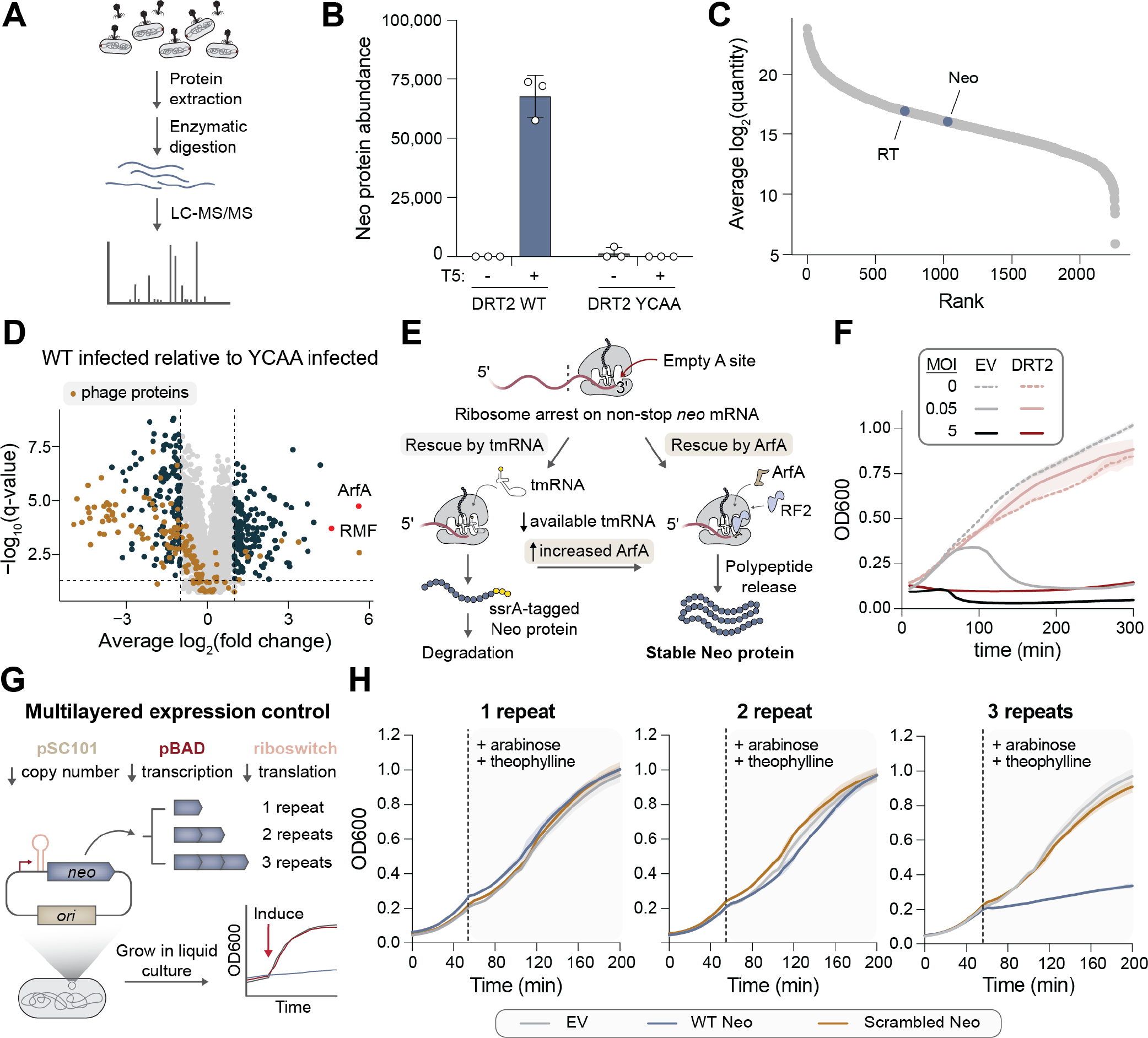
Neo proteins induce programmed cellular dormancy. (A) Schematic of experimental approach to detect Neo in phage-infected cells by liquid chromatography with tandem mass spectrometry (LC-MS/MS). **(B)** Bar graph quantifying Neo protein quantity from cells tested in the indicated conditions. Data are mean ± s.d. (n = 3). **(C)** Abundance of RT and Neo proteins relative to the *E. coli* proteome in phage-infected cells expressing WT DRT2. **(D)** Differential protein abundance in T5-infected cells expressing DRT2 WT or YCAA. Phage proteins are colored in brown, and ArfA and RMF are colored in red and labeled. All other differentially abundant proteins (fold change > 2 and FDR < 0.05) are colored in dark blue. **(E)** Schematic of alternative ribosome rescue pathway mediated by ArfA, which would release Neo proteins from ribosomes stalled on non-stop *neo* mRNAs without targeting them for degradation (right), unlike the tmRNA pathway (left). **(F)** Growth curves of strains transformed with empty vector (EV) or the WT DRT2 system, +/T5 phage at the indicated multiplicity of infection (MOI). Shaded regions indicate the standard deviation across independent biological replicates (n = 3). **(G)** Schematic of cloning and inducible expression strategy to monitor the physiological effects of Neo polypeptides of variable repeat length. **(H)** Growth curves of strains transformed with WT or scrambled Neo sequences of the indicated repeat lengths, alongside an empty vector (EV) control. The dashed line indicates the point of induction with arabinose and theophylline. Shaded regions indicate the standard deviation across independent biological replicates (n = 3).

To gain further insights into the physiological consequences of Neo expression, we performed additional MS-based proteomics experiments using a more standard trypsin-based digestion procedure, and analyzed the differential protein abundance between T5 phage-infected cells expressing WT or YCAA-mutant *Kpn*DRT2. Phage proteins were widely depleted in WT cells, as expected for a protective immune response (**Fig. 4D**). On the host side, two significantly enriched cellular factors immediately captured our attention — ArfA and RMF — due to their associations with ribosome stress and ribosome hibernation, respectively (**Fig. 4D**). ArfA (Alternative ribosome-rescue factor A) is a translation factor that specifically rescues ribosomes stalled on aberrant mRNAs lacking a stop codon, acting as an alternative to the tmRNA pathway that tags nascent polypeptide chains for degradation (*30*, *31*). ArfA is known to be specifically upregulated under conditions of tmRNA depletion (*32*), and its ribosome rescue activity in *neo*-expressing cells would elegantly resolve the conundrum of how stop codon-less *neo* mRNAs are nonetheless translated into functional proteins (**Fig. 4E**). Meanwhile, RMF (Ribosome modulation factor) is a ribosome-associated protein that directs the assembly of 70S ribosomes into inactive 100S dimers during stationary phase (*33*, *34*), and is activated by the alarmone ppGpp, a known trigger of growth arrest and cellular dormancy (*35*, *36*). Thus, the upregulation of ArfA supports the likely mechanism by which Neo polypeptides are translated, and the induction of RMF suggests that Neo production is associated with cellular dormancy.

Abortive infection and programmed dormancy have emerged in recent years as common mechanisms by which bacterial immune systems provide population-level immunity against phage infection, as host shutdown of metabolic processes prevents phage replication, and consequently, viral spread (*11*, *37*). To investigate whether DRT2 uses a similar immune mechanism, we performed phage infection assays in liquid culture at varying multiplicities of infection (MOI). DRT2-expressing cultures survived T5 phage infection at low MOI, but infection at high MOI led to stalling of growth (**Fig. 4F**). Further analysis of cultures infected at high MOI revealed that DRT2 effectively blocked T5 replication (**fig. S5B**), and that the growth-arrested cells remained viable (**fig. S5C**), altogether supporting a mechanism of phage defense via programmed dormancy.

We next sought to test the physiological effects of recombinant Neo expression, reasoning that the many intricate steps involved in Neo production from the ncRNA locus might add confounding factors to protein function analyses. We initially attempted to clone *neo* onto a standard inducible expression vector and test the hypothesis that *neo* expression would be sufficient to trigger cellular dormancy. Yet repeated attempts to clone expression vectors with more than 2 repeats of WT *neo* proved unsuccessful, compared to scrambled control sequences that could be cloned with high efficiency, and the few colonies that emerged consistently exhibited frameshift mutations or lacked the *neo* insert altogether (**fig. S5D,E**). These qualitative results suggested that Neo may potently arrest cell growth, and that its leaky expression had prevented the isolation of positive clones. Intriguingly, this effect was only observed with Neo repeat lengths of 3 or more.

To circumvent this challenge, we adopted an alternative strategy (*38*), in which *neo* genes were placed on a low-copy vector under the control of a tightly regulated pBAD promoter and theophylline riboswitch (**Fig. 4G**). This multilayered strategy for control of *neo* expression — which evokes the elaborate regulation of *neo* expression by native DRT2 loci — enabled the isolation of the desired clones. We then transformed cells with expression vectors encoding WT or scrambled *neo* with 1-3 repeats, and monitored cell growth in liquid culture before and after inducing *neo* expression with arabinose and theophylline. Strikingly, only the 3-repeat WT Neo construct exhibited any growth defect compared to an empty vector control (**Fig. 4H**). To assess whether the growth-arrested cells could recover from dormancy, we plated cells from the final time point of the liquid culture experiment on solid media supplemented with either repressor (glucose) or inducer (arabinose and theophylline). We found that cells expressing 3-repeat WT Neo exhibited a ∼10^2^-fold increase in colony-forming units when plated on repressor versus inducer (**fig. S5F**), indicating strong recovery from Neo-induced dormancy.

Considered together, these results suggest that the intricate gene synthesis mechanism encoded by *Kpn*DRT2 may have evolved in order to strictly control the production of an effector protein whose toxicity is too potent to be safely controlled by conventional regulatory strategies.

### *Neo* gene synthesis and Neo protein toxicity is a broadly conserved phage defense strategy

Equipped with a wealth of mechanistic information on the production of Neo protein by *Kpn*DRT2, we set out to explore the evolutionary conservation of this gene synthesis strategy for antiviral defense. Starting with a large phylogenetic tree of DRT2 homologs (**fig. S6A and Supplementary Table S1**), we used covariance models to annotate *RT*-associated ncRNAs and then extracted the putative *neo* gene and Neo protein sequence based on the expected mechanism of template jumping and absence of in-frame stop codons (**Fig. 5A and Materials and Methods**). Our pipeline identified candidate ncRNAs and Neo proteins for the vast majority of DRT2 systems that were related to *Kpn*DRT2 (**Fig. 5B**), revealing broad conservation of this unique mechanism for concatenated gene synthesis. Notably, sequence motifs expected to be critical for *neo* gene synthesis and expression, including ACA-1, ACA-2, and repeat junction-flanking promoter elements, were also strongly conserved across diverse homologs (**fig. S6B**). Iterative generation of additional covariance models also enabled ncRNA prediction for more divergent DRT2 clades, but Neo protein annotation was more challenging, suggesting the possibility of alternative mechanisms of RCRT and *neo* gene expression (**fig. S6A,C**).

**Fig. 5.**
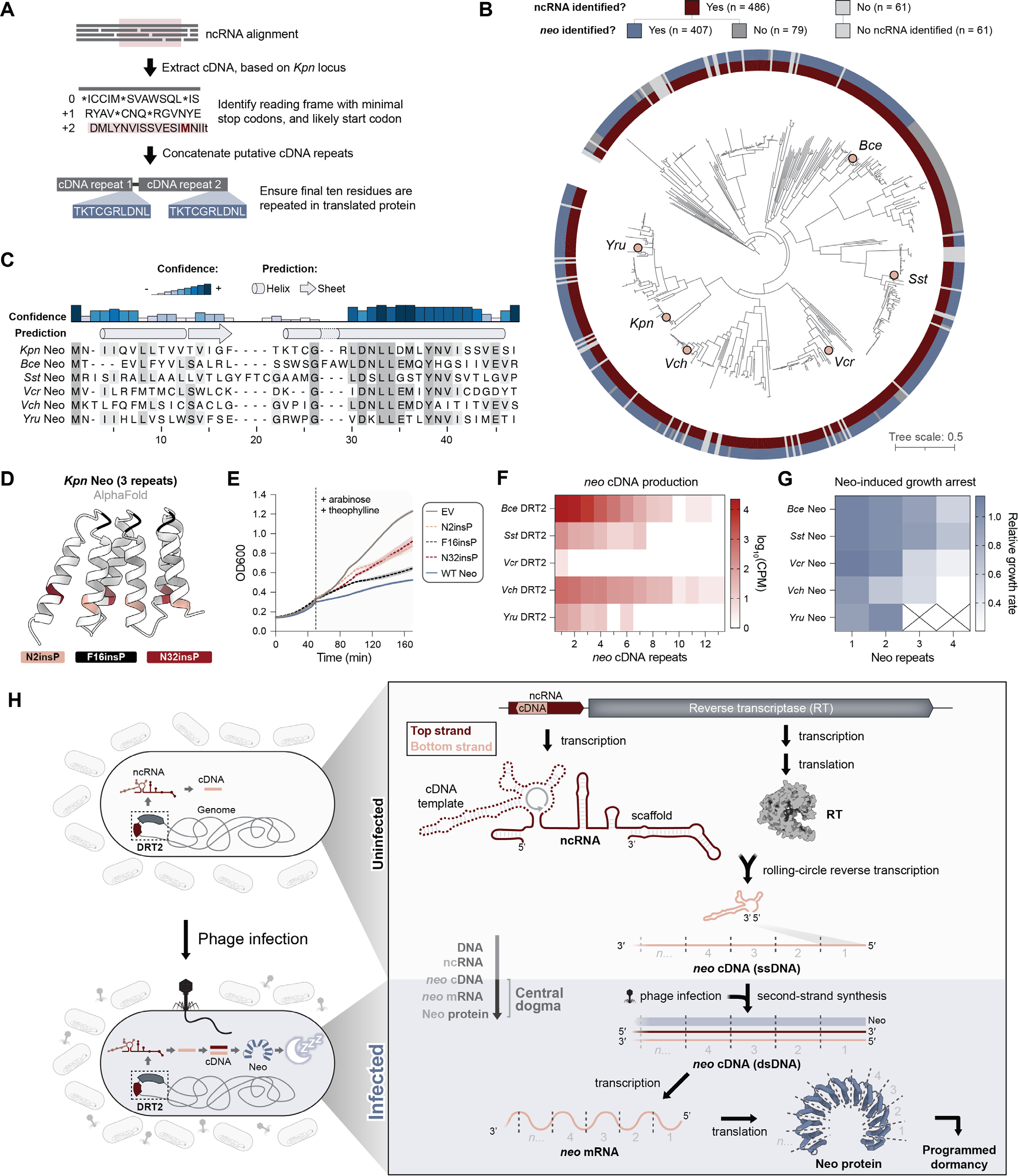
Concatenated *neo* genes and programmed dormancy are a broadly conserved phage defense mechanism. (A) Schematic for the automated detection of putative Neo proteins in homologous DRT2 operons. **(B)** Phylogenetic tree of *Kpn*DRT2 homologs, with outer rings showing the widespread presence of RT-associated ncRNAs and putative Neo proteins. Homologs selected for experimental testing are indicated with pink circles. **(C)** Multiple sequence alignment (MSA) and secondary structure prediction of Neo proteins identified in **B**. A single Neo repeat is shown for all homologs; shading indicates amino acid conservation. **(D)** AlphaFold prediction of a 3-repeat Neo polypeptide, showing the sites of proline mutagenesis tested in **E**. Prolines were inserted C-terminal to the indicated residues within each of 3 concatenated repeats. **(E)** Growth curves of strains transformed with 3-repeat Neo constructs containing the indicated proline insertions, alongside an empty vector (EV) control. The dashed line indicates the point of induction with arabinose and theophylline. Shaded regions indicate the standard deviation across independent biological replicates (n = 3). **(F)** Heat map showing the distribution of *neo* cDNA repeat lengths in cells expressing the indicated DRT2 homologs. Data are plotted as log_10_(CPM) from Nanopore sequencing of total DNA. **(G)** Heat map showing the growth rates of cells expressing Neo homologs with the indicated repeat lengths. Growth rates are normalized to an EV control and represent the mean of independent biological replicates (n = 3). Empty cells with X indicate Neo expression constructs that could not be successfully cloned, presumably due to toxicity. **(H)** Model for the antiphage defense mechanism of DRT2 systems. RT enzymes bind the scaffold portion of associated ncRNAs and produce concatenated cDNA products via rolling-circle reverse transcription (RCRT). Phage infection triggers second-strand synthesis, yielding a dsDNA molecule that is transcribed into never-ending ORF (*neo*) mRNAs. *Neo* translation exploits a ribosome rescue pathway to produce Neo proteins that potently arrest cell growth, protecting the larger bacterial population from the spread of phage.

Next, we investigated the amino acid sequence of diverse Neo proteins in more detail. Bioinformatics analyses of multiple sequence alignments failed to identify any functional domains or similarities to known proteins, but they did reveal high-confidence predictions of α-helical secondary structural elements (**Fig. 5C**). Using a 3-repeat Neo sequence, we predicted the 3D protein fold using multiple independent methods, which yielded a structure reminiscent of HEAT repeats and other alpha solenoids consisting of repeating antiparallel α-helices (*39*, *40*) (**Fig. 5D and fig. S6D**). To test this model, we introduced helix-breaking proline residues into either the loop connecting two α-helices, or into the helices directly (**Fig. 5D**), and assessed the effects of these perturbations on cell growth. Consistent with our structural model, insertions into either helix eliminated Neo-induced growth arrest, whereas the loop insertion mutant exhibited a dormancy phenotype similar to the WT Neo sequence (**Fig. 5E**).

Having found that Neo proteins exhibit conserved α-helical folds, we sought to experimentally test the conservation of additional critical features of Neo production and cellular function. We selected and cloned 5 diverse DRT2 homologs (**Fig. 5B and fig. S7A**), and performed Nanopore sequencing of total DNA from cells transformed with DRT2 expression vectors to assess the distribution of *neo* cDNA repeat lengths. Remarkably, nearly all of the tested systems exhibited RCRT upon heterologous expression in *E. coli*, and the cDNAs spanned a wide range of abundance levels and repeat lengths (**Fig. 5F**). We also tested the effect of recombinant expression of the Neo proteins predicted to be encoded by these concatenated cDNAs. In all cases, Neo homolog expression led to repeat length-dependent growth arrest, further confirming the requirement for cDNA repeat concatenation in the programmed dormancy mechanism to defend against phage infection (**Fig. 5G and fig. S7B**).

Collectively, these experiments establish the generalizability of the *Kpn*DRT2 gene synthesis and antiphage defense mechanisms across a large swath of related immune systems.

## DISCUSSION

Our work reveals an unprecedented mechanism of antiviral immunity mediated by DRT2 defense systems (**Fig. 5H)**. In uninfected cells, the ncRNA and RT enzyme are constitutively expressed from a single promoter, leading to synthesis of a repetitive single-stranded cDNA via precise, programmed template jumping that mediates RCRT. Upon phage infection, second-strand synthesis is triggered, leading to the accumulation of double-stranded, concatenated cDNA molecules. A promoter created across the junction between adjacent cDNA repeats then leads to abundant expression of heterogeneously sized mRNA encoding a stop codon-less, never-ending ORF (*neo*). It is the production of Neo protein, we propose, that acts as the effector arm of the immune system by rapidly arresting cell growth and inducing programmed dormancy, thus protecting the larger bacterial population from the spread of phage.

This pathway complicates textbook descriptions of the central dogma of molecular biology by highlighting complex and repeated transitions back and forth between DNA- and RNA-based carriers of genetic information, before translation finally yields a protein product. Furthermore, it challenges the universal paradigm that genes are encoded linearly along the chromosomal axis. Genes across all three domains of life are arranged in a polarized and singular orientation from head to tail, even considering the existence of intron splicing and discontinuous exon joining in eukaryotes. Although *neo* proto-genes are similarly arranged, synthesis of the mature gene form requires RNA-templated concatenation of the tail of one proto-gene to the head of another. When considered alongside other examples of strategies used by mobile elements to compactly encode genetic information, including ribosomal frameshifting, overlapping ORFs, and nested genes, our work adds another layer of complexity to the ways in which protein-coding sequences can be stored in the genome.

Why would such an elaborate immune pathway and gene synthesis mechanism have evolved? One potential explanation is the need for stringent control of Neo expression. Our initial attempts to clone recombinant Neo for functional testing were fraught with technical challenges, including the rapid selection of loss-of-function mutants due to cellular toxicity, suggesting an extreme fitness cost associated with even low levels of Neo expression. It is likely that genomic encoding of pre-assembled *neo* genes, under standard transcriptional control mechanisms, would pose intolerable autoimmune risk to the host. We therefore hypothesize that gating Neo expression behind multiple layers of regulation, including RNA-templated cDNA synthesis, phage-triggered dsDNA synthesis, and ArfA-mediated ribosome rescue, likely allowed a potent dormancy factor to be stably maintained as part of the immune response.

Perhaps the most mysterious aspect of DRT2 immunity that remains to be elucidated is the structure and molecular function of Neo. Although we detected unambiguous Neo-derived peptides in phage-infected cells via mass spectrometry, the necessary protease digestion steps precluded determination of its size. Our results indicate that a Neo polypeptide of at least 3 repeats in length is necessary and sufficient to induce cell dormancy, but it is worth noting that *neo* mRNAs range in size from 200 to >5,000 nt, and that translation could in theory initiate from within any *neo* repeat, each of which contains its own RBS. Thus, we anticipate that native Neo proteins are similarly heterogeneous in size, potentially spanning hundreds to thousands of amino acids. Mass spectrometry-based proteomics also highlighted ribosome hibernation as a downstream effect of DRT2 immune system function, though additional experiments will be necessary to determine whether RMF activation is a direct of Neo, or, more likely, an indirect consequence of programmed dormancy. The fact that RMF activation is driven by the alarmone ppGpp (*35*), which causes dormancy and growth arrest and is itself synthesized on ribosomes (*41*), raises the possibility that Neo directly induces this cellular pathway. Finally, the conservation of α-helical repeat (αRep) domains fused to reverse transcriptase domains in Class 1 DRT systems (*19*), which strongly resemble the predicted α-helical fold of Neo, tantalizingly suggest a potential unifying theme for the effector functions across all DRT systems.

Our discovery of highly efficient RCRT activity represents a unique biochemical behavior that produces concatenated, repetitive cDNA molecules with precise junction sequences. Although many other characterized reverse transcriptases exhibit intermolecular template switching activity in vitro (*42*, *43*), the intramolecular template jumping mechanism we describe here is the first report of such an activity creating de novo protein-coding genes *in vivo*, with direct implications for biological function. Additional work will be needed to determine the specific adaptations in the RT enzyme that facilitate this activity, but our bioinformatics analyses and experimental results point to the critical importance of conserved ncRNA structure and sequence motifs — in particular, the ACA motifs found abutting and within SL2. These motifs likely guide reannealing of the nascent cDNA transcript after one round of cDNA synthesis to a second template immediately upstream of the cDNA start site, thereby initiating a new round of cDNA synthesis. When considered alongside other well-studied examples of programmed template switching — such as the synthesis of subgenomic RNAs by coronaviral RNA-dependent RNA polymerases (*44*, *45*), and the synthesis of full-length genomic cDNAs by retroviral reverse transcriptases (*46*) — our work expands the diversity of products that can be generated by a single polymerase enzyme from its substrate. Furthermore, DRT2 represents the first biological system in which a reverse transcriptase natively performs RCRT. Intriguingly, it does so using a template that is not a closed circle, and thus differs from classic examples of rolling circle amplification associated with plasmid, phage, and viroid replication (*26*). These unique properties of DRT2-encoded enzymes offer considerable potential for biotechnology applications that leverage templated DNA production in vivo (*47–49*), but with the added advantage of programmed amplification. Notably, our data demonstrate that RCRT is maintained with mutation of SL5 or the reverse transcription start site, suggesting that DRT2 could be harnessed to produce high-copy concatenated cDNAs with user-defined sequences.

The identification of Neo proteins disrupts conventionally held notions of features that define protein coding genes, as well as our broader understanding of genome composition. Current genome annotations algorithms generally rely on the definition of an ORF as a translated sequence of at least 30 amino acids bounded by a start and stop codon (*50*). However, only 26 codons of *neo* are identifiable from a linear view of the *K. pneumoniae* genome, and the proto-gene lacks a stop codon. Thus, *neo* genes are hidden in regions of genomes previously thought to be exclusively non-coding, suggesting that alternative bioinformatics approaches will be needed in order to discover similar genes that elude standard methods of ORF prediction. These findings seem especially important when considering the large proportion of non-coding DNA in higher eukaryotes. For example, only ∼1.5% of the human genome is thought to encode proteins (*51*). While much of the remaining ∼98.5% of genome content encodes RNAs with known or predicted gene regulatory functions, we posit that additional examples of Neo-like, non-canonical protein coding genes likely await future discovery in our own genomes.

## ACKNOWLEDGEMENTS

We thank S. Pesari and Z. Akhtar for laboratory support, A. Bernheim and J. Bondy-Denomy for helpful guidance on phage infection experiments, E. Semenova for helpful advice on ssDNA sequencing library preparation, A.W.P. Fitzpatrick for insightful discussions on Neo function, B. Wiedenheft for suggesting the Neo acronym, M. Laub for generously providing T5 phage, L.F. Landweber for qPCR instrument access, and the JP Sulzberger Columbia Genome Center for NGS support.

## AUTHOR CONTRIBUTIONS

S.T. and S.H.S. conceived of and designed the project. S.T. performed most experiments and experimental analyses. V.C. performed cell growth and plaque assays, time course experiments, and helped with cloning. D.J.Z. performed initial phylogenetic analyses and selected DRT2 homologs, generated the ncRNA covariance model, performed cell growth and plaque assays, and helped with cloning. R.Z. performed RT-qPCR and cell growth assays, and helped with cloning. G.D.L. performed and helped with the analysis of Nanopore sequencing experiments. T.W. performed phylogenetic analyses and bioinformatically identified Neo homologs. L.C.T. prepared samples and performed mass spectrometry experiments. M.W. performed cell growth assays. M.W.G.W. helped with the design and analysis of Neo protein expression data. J.T.G. contributed to the interpretation of rolling-circle reverse transcription results. L.E.B. advised on the design and interpretation of Northern blotting experiments. M.J. advised on the design and interpretation of mass spectrometry experiments. S.T. and S.H.S. discussed the data and wrote the manuscript, with input from all authors.

## DATA AND MATERIALS AVAILABILITY

Next-generation sequencing data will be made available in the National Center for Biotechnology Information (NCBI) Sequence Read Archive upon publication. Additional datasets generated and analyzed in the current study are available from the corresponding author upon reasonable request.

## SUPPLEMENTARY MATERIALS

Materials and Methods Figs. S1 to S7

Tables S1 to S5 (provided separately)

