## Supplementary Materials for "De novo gene synthesis by an antiviral reverse transcriptase"

**The PDF file includes:**

Materials and Methods
Figs. S1 to S7

**Other Supplementary Materials for this manuscript include the following:**

Tables S1 to S5

### MATERIALS AND METHODS

Plasmid and *E. coli* strain construction:

All strains and plasmids used in this study are described in **Supplementary Tables S2** and **S3**, respectively. Briefly, plasmids were cloned using a combination of methods, including Gibson assembly, restriction digestion-ligation, ligation of hybridized oligonucleotides, Golden Gate Assembly, and around-the-horn PCR. Plasmids were cloned and propagated in *E. coli* strain NEB Turbo (sSL0410), and all experiments were performed in *E. coli* str. K-12 substr. MG1655 (sSL0810). Clones were verified by Sanger sequencing or whole plasmid sequencing. pLG007 (Ec86 retron) and pLG010 (DRT type 2) were gifts from Feng Zhang (Addgene plasmids # 157885, # 157888) (*13*).

Phage amplification and plaque assays:

Phage T5 (a gift from Michael Laub) was amplified in liquid culture by diluting an overnight culture of MG1655 cells 1:100 in 10 mL fresh LB media, adding 50 uL of phage, and incubating at 37 °C for 3-4 hours. Chloroform was added to a final concentration of 5% to facilitate complete bacterial lysis, after which the lysate was centrifuged at 4,000 x *g* for 10 min to pellet cell debris. The supernatant was passed through a sterile 0.22 µm filter, and the phage-containing filtrate was stored at 4 °C.

Small-drop plaque assays were performed as follows: *E. coli* str. K-12 substr. MG1655 (sSL0810) was transformed with the indicated plasmid construct (see **Supplementary Table S3** for plasmid descriptions and sequences) and plated on solid LB media. Single colonies were inoculated in liquid LB media containing the appropriate antibiotic and grown overnight at 37 °C with shaking. The next day, 100 µL of overnight culture were mixed with 4 mL freshly prepared molten soft agar (0.5% agar in LB media containing the appropriate antibiotic) at 42 °C and poured over solid bottom agar (1.5% agar in LB media containing the appropriate antibiotic) in a 10 cm Petri dish. The soft agar was allowed to solidify for 15 min at RT, during which 10× serial dilutions of phage T5 in LB were prepared. For plating, 3 µL of each phage dilution were spotted onto the surface of the soft agar lawn and were allowed to dry uncovered for 10 min under a laminar flow hood. Plates were incubated at 37 °C for 8-16 hours to allow the formation of plaques. After selecting a phage dilution with clearly distinguishable plaques, plaque forming units (PFU) $mL^{-1}$ were calculated using the following formula: . Phage defense activity was assessed by calculating the fold reduction in efficiency of plating (EOP), which was determined by dividing the PFU $mL^{-1}$ obtained on a lawn of empty vector (EV) control cells by the PFU $mL^{-1}$ obtained on a lawn of defense system-expressing cells.

RNA and cDNA immunoprecipitation and sequencing (RIP-seq and cDIP-seq):

*E. coli* str. K-12 substr. MG1655 (sSL0810) was transformed with plasmids encoding C-terminally 3×FLAG-tagged Retron-Eco1 or N-terminally 3×FLAG-tagged *Kpn*DRT2 (WT or RT-inactive YCAA mutant), as well as their native flanking sequences (see **Supplementary Table S3** for plasmid sequences). Individual colonies were inoculated in liquid LB with chloramphenicol (25 µg $mL^{-1}$) and grown at 37 °C to $OD_{600}$ of 0.5. For experiments +/- phage infection, 40 mL cultures were split in half, and phage T5 was added to one half at a multiplicity of infection (MOI) of 5, which was calculated as the ratio of phage PFU to bacterial colony forming units (CFU), assuming $8\times10^8$ CFU in 1 mL culture at $OD_{600}$ of 1.0. Uninfected and infected cultures were grown for 1 hr at 37 °C. For experiments without phage infection, 20 mL cultures were grown to $OD_{600}$ of 0.5 and directly harvested. Cells were harvested by centrifugation at 4,000 x *g* for 10 min at 4 °C, and the supernatant was removed. The pellet was washed with 5 mL of cold TBS

(20 mM Tris-HCl, pH 7.5 at 25 °C, 150 mM NaCl) and spun down again as before. The supernatant was removed, and the pellet was washed with 1 mL of cold TBS before centrifugation at 10,000 x *g* for 5 min at 4 °C. The supernatant was removed, and the pellet was flash-frozen in liquid nitrogen and stored at -80 °C.

Antibodies for immunoprecipitation were conjugated to magnetic beads as follows: for each sample, 60 µL Dynabeads Protein G (Thermo Fisher Scientific) were washed 3× in 1 mL IP lysis buffer (20 mM Tris-HCl, pH 7.5 at 25 °C, 150 mM KCl, 1 mM $MgCl_2$, 0.2% Triton X-100), resuspended in 1 mL IP lysis buffer, combined with 20 µL anti-FLAG M2 antibody (Sigma-Aldrich, F3165), and rotated for > 3 hours at 4 °C. Antibody-bead complexes were washed 2× to remove unconjugated antibodies and resuspended in 60 µL of IP lysis buffer per sample.

Flash-frozen pellets were thawed on ice and resuspended in 1.2 mL IP lysis buffer supplemented with 1× cOmplete Protease Inhibitor Cocktail (Roche) and 0.1 U $\mu L^{-1}$ SUPERase•In RNase Inhibitor (Thermo Fisher Scientific). To lyse cells, samples were sonicated using a 1/8″ sonicator probe for 1.5 min total (2 s ON, 5 s OFF) at 20% amplitude. To clear cell debris and insoluble material, lysates were centrifuged at 21,000 x *g* for 15 min at 4 °C, and 1 mL supernatant was transferred to a new tube. At this point, two small volumes of each sample (10 µL for RIP-seq and 10 µL for cDIP-seq) were set aside as "input" starting material and stored at -80 °C. For immunoprecipitation, each sample was combined with 60 µL antibody-bead complex and rotated overnight at 4 °C. The next day, each sample was washed 3× with 1 mL ice-cold IP wash buffer (20 mM Tris-HCl, pH 7.5 at 25 °C, 150 mM KCl, 1 mM $MgCl_2$), using a magnetic rack to immobilize the beads in between each wash. During the final wash, each sample was separated into two separate 500 µL volumes for downstream RIP or cDIP processing.

For RIP elution, the supernatant was removed, and beads were resuspended in 750 µL TRIzol (Thermo Fisher Scientific). After 5 min incubation at RT, the supernatant containing eluted RNA was transferred to a new tube and combined with 150 µL chloroform. Samples were mixed vigorously by inversion, incubated at RT for 3 min, and centrifuged at 12,000 x *g* for 15 min at 4 °C. RNA was isolated from the upper aqueous phase using the RNA Clean & Concentrator-5 kit (Zymo Research) and eluted in 15 µL RNase-free water. RNA from input samples was isolated in the same manner using TRIzol and column purification. Purified RNA was stored at -80 °C before proceeding to library preparation.

For cDIP elution, the supernatant was removed, and beads were resuspended in 90 µL IP wash buffer and treated with 5 µg RNase A (Thermo Fisher Scientific) for 30 min at 37 °C. Input samples were adjusted to 90 µL with IP wash buffer and treated with RNase A in parallel. SDS was added to IP and input samples to a final concentration of 1%, and samples were treated with 25 µg Proteinase K (Thermo Fisher Scientific) for 30 min at 55 °C. Beads were immobilized using a magnetic rack, and the supernatant containing eluted DNA was transferred to a new tube. DNA was isolated using the Monarch PCR and DNA Cleanup kit (NEB), following the Oligonucleotide Cleanup protocol and eluting in 15 µL DNase-free water. For Retron-Eco1 samples, DNA was treated with DBR1 (Origene) in reactions containing 2 µL DNA, 0.5 µL DBR1, 1× rCutSmart in 10 µL total volume, in order to cleave the 2′-5′ phosphodiester linkage between msRNA and msDNA. Reactions were cleaned up using the Monarch PCR and DNA Cleanup kit (NEB), with elution in 15 µL DNase-free water. Purified DNA was stored at -80 °C before proceeding to library preparation.

For RIP-seq library preparation (input and RIP eluates), RNA was fragmented by random hydrolysis by combining 7 µL RNA, 6 µL water, and 2 µL NEBuffer 2, and heating to 92 °C for 2 min. To remove DNA and prepare RNA ends for adapter ligation, samples were treated with 2 µL TURBO DNase (Thermo Fisher Scientific) and 2 µL RppH (NEB) in the presence of 1 µL SUPERase•In RNase Inhibitor for 30 min at 37 °C. This was followed by treatment with 1 µL T4 PNK (NEB) in 1× T4 DNA ligase buffer (NEB) for 30 min at 37 °C. Reactions were column-purified using the Zymo RNA Clean & Concentrator-5 kit and eluted in 10.5 µL RNase-free water. RNA concentrations were quantified using the DeNovix RNA Assay. Illumina sequencing libraries were prepared using the NEBNext Small RNA Library Prep kit, and libraries were sequenced on an Illumina NextSeq 500 in paired-end mode with 150 cycles per end.

For cDIP-seq library preparation, 2 µL of each input sample and 10 µL of each IP eluate were diluted to 15 µL with DNase-free water. Samples were denatured by heating at 95 °C for 2 min, and then immediately placed on ice. Ligation of Illumina adapters and conversion of ssDNA to dsDNA were performed using the xGen ssDNA & Low-Input DNA Library Prep Kit (IDT), and libraries were sequenced on an Illumina NextSeq 500 in paired-end mode with 150 cycles per end.

#### RIP-seq and total RNA-seq analyses:

RIP-seq and corresponding input datasets were processed using cutadapt (*52*) (v4.2) to remove Illumina adapter sequences, trim low-quality ends from reads, and filter out reads shorter than 15 bp. Reads were mapped to combined reference files containing the MG1655 genome (NC_000913.3) and relevant plasmid sequence, as well as the T5 genome (NC_005859.1) for +/- infection experiments, using bwa-mem2 (*53*) (v2.2.1) with default parameters. SAMtools (*54*) (v1.17) was used to sort and index alignments. Coverage tracks were generated using bamCoverage (*55*) (v3.5.1) with a bin size of 1, separation of top and bottom strand alignments, and scaling of coverage according to sequencing depth (based on the total number of reads passing initial trimming and length filtering). Coverage tracks were visualized in IGV (*56*).

For transcriptome-wide analyses of RNAs enriched by RIP-seq, aligned reads were assigned to annotated transcriptome features using featureCounts (*57*) (v2.0.2) with -s 1 for strandedness. The resulting counts matrices were passed to DESeq2 (*58*) to calculate fold-change and FDR (using the Benjamini-Hochberg procedure) between input and IP for each annotated transcript. Comparisons were visualized using ggplot2, plotting the "baseMean" (mean normalized counts across all conditions) against $\log_2$(fold change). All comparisons included three independent biological replicates.

For counting of *neo* repeat junction-spanning reads in RIP input (i.e., total RNA) samples, a custom reference sequence was made which consisted of two concatenated *neo* cDNA repeats. A 20-bp feature annotation was added, centered at the repeat–repeat junction. Reads were aligned to the custom reference sequence using bwa-mem2, and featureCounts was used to count alignments spanning the junction annotation. The resulting counts were normalized for sequencing depth.

#### cDIP-seq and total DNA sequencing analyses:

Adapter trimming, quality trimming, and length filtering of cDIP-seq and corresponding input datasets were performed as described above for RIP-seq experiments. Trimmed and filtered reads were mapped to combined reference files, sorted, indexed, and plotted onto coverage tracks as described above. Alignments over annotated transcriptome features were counted using featureCounts with -s 2 for strandedness. The resulting counts matrices were processed by DESeq2 and plotted as described above. All transcriptome-wide comparisons were performed using three independent biological replicates.

In order to plot cDNA 5′ and 3′ ends over the *Kpn*DRT2 ncRNA locus, cDIP-seq alignment coordinates were extracted using the bamtobed utility from bedtools (*59*) (v2.31.0). The 5′ boundary of each read pair was determined as the start coordinate of read 1, for transcripts on the top strand, or the end coordinate of read 1, for transcripts on the bottom strand. Meanwhile, the 3′ boundary of each read pair was determined as the end coordinate of read 2, for transcripts on the top strand, or the start coordinate of read 2, for transcripts on the bottom strand. The boundary coordinates thus defined for each read pair were plotted as a histogram over the *Kpn*DRT2 ncRNA locus.

For counting of reads mapping to the *Kpn*DRT2 cDNA, a custom annotation file was created which defined the DRT2 cDNA feature based on the coverage boundaries from cDIP-seq of *Kpn*DRT2. Alignments over this feature were counted using featureCounts with -s 2 for strandedness and --minOverlap 60. Counting of *neo* repeat–repeat junction-spanning reads was performed as described above for RIP input samples. The proportion of junction-spanning versus non-junction-spanning cDNA alignments was calculated by dividing the junction-spanning read counts by the total number of reads mapped to the custom concatenated reference sequence.

To analyze cDIP-seq reads with soft-clipped extensions beyond the DRT2 cDNA coverage boundary, cutadapt was used to extract reads containing the full-length *Kpn*DRT2 cDNA and then trim the cDNA repeat sequence from the 5′ end of the read. This step produced trimmed reads containing only the portion of the read extending beyond the coverage boundary. A custom bash script was used to calculate the lengths of the extensions. The extensions were subsequently mapped back to the combined MG1655 genome, T5 phage, and DRT2 plasmid reference using bwa-mem2. Coverage tracks of the alignments were generated using bamCoverage.

<u>dRNA-seq</u>:

To precisely map the transcription start site of the *Kpn*DRT2 ncRNA, a custom RNA-seq library preparation protocol was used to enrich primary transcripts from the total RNA pool, as previously described (*22*). *E. coli* MG1655 cells transformed with a plasmid encoding *Kpn*DRT2 were grown to exponential phase, and total RNA was extracted using TRIzol (Thermo Fisher Scientific). 1 µg of total RNA was fragmented in 1× NEBuffer 2 by heating at 92 °C for 1.5 min. DNase treatment was performed with 1 µL TURBO DNase (Thermo Fisher Scientific) in the presence of 1 µL SUPERase•In RNase Inhibitor (Thermo Fisher Scientific) for 10 min at 37 °C. Samples were treated with 1 µL T4 PNK in 1× T4 DNA ligase buffer (NEB) at 37 °C for 30 min and purified using the Zymo RNA Clean & Concentrator-5 kit. To enrich primary transcripts with tri-phosphorylated 5′ ends, samples were treated with 1 µL of Terminator Exonuclease (Biosearch Technologies) in 1× Terminator Reaction Buffer A (Biosearch Technologies) supplemented with 0.5 µL SUPERase•In RNase Inhibitor (Thermo Fisher Scientific). Reactions were incubated at 30 °C for 1 hour and stopped by adding EDTA to a final concentration of 5 mM. Samples were purified using the Zymo RNA Clean & Concentrator-5 kit, and then treated with 2 µL RppH (NEB) in 1× NEBuffer 2 supplemented with 1 µL SUPERase•In RNase Inhibitor (Thermo Fisher Scientific). Reactions were incubated at 37 °C for 30 min and purified using the Zymo RNA Clean & Concentrator-5 kit. Illumina sequencing libraries were prepared using the NEBNext Small RNA Library Prep kit, and libraries were sequenced on an Illumina NextSeq 500 in single-end mode with 75 cycles per end.

Adapter trimming, quality trimming, and length filtering of dRNA-seq reads were performed as described above for RIP-seq experiments. Trimmed and filtered reads were mapped to reference files using bowtie2 (*60*) (v2.4.5) with default parameters. Alignments were sorted and indexed as described above. The locations of RNA 5′ ends over the *Kpn*DRT2 ncRNA locus were determined and plotted as described above for cDNA 5′ end analysis.

<u>Term-seq:</u>

Term-seq was performed to enrich the 3′ ends of transcripts, as previously described (*23*), using the same RNA sample as used for dRNA-seq. 1 µg of total RNA was treated with 1 µL TURBO DNase in 1× TURBO DNase Buffer (Thermo Fisher Scientific) supplemented with 1 µL SUPERase•In RNase

Inhibitor (Thermo Fisher Scientific) for 10 min at 37 °C, followed by cleanup using the Zymo RNA Clean & Concentrator-5 kit. Ligation of an i7 Illumina adapter to RNA 3′ ends was performed using the NEB-Next Small RNA Library Prep kit, followed by cleanup using the Zymo RNA Clean & Concentrator-5 kit. Samples were fragmented in 1× NEBuffer 2 by heating at 92 °C for 1.5 min, then treated with 2 μL RppH (NEB) in the presence of 1 μL SUPERase•In RNase Inhibitor (Thermo Fisher Scientific) for 30 min at 37 °C. This was followed by treatment with 1 μL T4 PNK in 1× T4 DNA ligase buffer (NEB) at 37 °C for 30 min and cleanup using the Zymo RNA Clean & Concentrator-5 kit. Illumina library preparation continued with the remainder of the NEBNext Small RNA Library Prep protocol after the initial i7 adapter ligation step. Libraries were sequenced on an Illumina NextSeq 500 in single-end mode with 75 cycles per end.

Adapter trimming, quality trimming, and length filtering of Term-seq reads were performed as described above for RIP-seq experiments. Trimmed and filtered reads were mapped to reference files using bowtie2 (*60*) (v2.4.5) with default parameters. Alignments were sorted and indexed as described above. The locations of RNA 3′ ends over the *Kpn*DRT2 ncRNA locus were determined and plotted as described above for cDNA 3′ end analysis.

<u>Long-read DNA sequencing</u>:

Total DNA was extracted from *E. coli* str. K-12 substr. MG1655 (sSL0810) cells transformed with the indicated DRT2 expression vectors, using the Wizard Genomic DNA purification kit (Promega). For experiments performed in the absence of phage infection, single-stranded DNA was converted to double-stranded DNA using the Adaptase and Extension modules of the xGen ssDNA & Low-Input DNA Library Prep Kit (IDT). DNA was then purified using 1.2× AMPure XP beads (Beckman Coulter). This dsDNA conversion step was omitted for experiments performed in the presence of phage, as the Adaptase reaction is biased toward short ssDNA fragments (see user manual), and because phage infection is expected to trigger the *in vivo* conversion of single-stranded DRT2 cDNA to double-stranded DNA. DNA samples were prepared for long-read sequencing using the Native Barcoding Kit (Oxford Nanopore), following the manufacturer's protocol. Sequencing using an ONT MinION was performed with real time basecalling, barcode balancing, minimum read length of 200 bp, read splitting on, and minimum Q score of 8.

Adapter trimming and barcode trimming were performed with guppy barcoder (v6.5.7). To filter out non-cDNA reads, minimap2 (*61*) (v2.26) was used to align reads to plasmid reference sequences in which the expected cDNA region had been removed, as well as to the *E. coli* genome. Unmapped reads were then extracted for downstream analysis using SAMtools. A custom script was used to quantify the number of cDNA repeats detected in each sequencing read, and counts were normalized to the total number of sequenced reads for each sample. For visualization of concatenated cDNAs from the phage-infected *Kpn*DRT2 sample, reads were aligned to an artificial reference sequence using the built-in aligner in Geneious with medium sensitivity and an iteration of up to five times. The artificial reference sequence was created by concatenating up to 50 repeats of the cDNA template. To ensure that reads were aligned to the start of the cDNA concatemer, and not stochastically across the repeated sequence, an 'anchor' sequence was appended to the 5′ end of the first strand in all filtered sequences and the beginning of the cDNA concatemer sequence, thereby enforcing synchronous alignment starting at the 5′ end of the cDNA. Coverage over the reference cDNA concatemer was then exported for visualization.

Liquid chromatography with tandem mass spectrometry:

*E. coli* str. K-12 substr. MG1655 (sSL0810) cells transformed with the indicated DRT2 expression vectors were grown at 37 °C in 50 mL LB with chloramphenicol (25 $\mu g\,mL^{-1}$) to $OD_{600}$ of 0.5. Phage T5 was added at MOI 5 and cultures were infected for 1 hour. Cells were harvested by centrifugation at 4,000 x *g* for 10 min at 4 °C, and the supernatant was discarded. The pellet was washed with 5 mL cold TBS (20 mM Tris-HCl, pH 7.5 at 25 °C, 150 mM NaCl) and spun down again as before. The supernatant was removed, and the pellet was washed with 1 mL of cold TBS before centrifugation at 20,000 x *g* for 5 min at 4 °C. The supernatant was removed, and the pellet was flash-frozen in liquid nitrogen and stored at -80 °C.

Flash-frozen pellets were thawed on ice and resuspended in 1 mL lysis buffer (100 mM ammonium bicarbonate, 2% sodium deoxycholate). Cells were sonicated using a 1/8″ sonicator probe for 1.5 min total (5 s ON, 10 s OFF) at 20% amplitude. Lysates were heated to 95 °C for 10 min. Protein concentrations were assessed using the Pierce BCA assay (Thermo Fisher Scientific). 50 µg of each sample were subjected to reduction by DTT and alkylation by IAA before being precipitated onto SP3 beads as previously described (*62*). The beads were washed and then the samples were split in two, to be digested under different digestion conditions. In one condition, proteins underwent on-bead digestion by trypsin, glu-c, and chymotrypsin; this protease mixture was specifically chosen to generate peptides from the *Kpn* Neo protein in an amino acid length range suitable for detection by LC-MS/MS (**fig. S5A**). In the other condition, proteins underwent on-bead digestion by trypsin alone. This more conventional digestion approach was adopted to facilitate the analysis of global proteomic changes that occurred under the different experimental conditions. Each of the proteases was added in a 1:50 enzyme:substrate ratio for overnight digestion at room temperature.

Whole proteome, label-free MS analyses were performed by data-independent acquisition (DIA). Approximately 1 µg of total peptides was analyzed on a Waters M-Class UPLC using a 15 cm IonOpticks Aurora Elite column (75 µm inner diameter; 1.7 µm particle size; heated to 45°C) coupled to a benchtop Thermo Fisher Scientific Orbitrap Q Exactive HF mass spectrometer. Peptides were separated at a flow rate of 400 nL/min with a 150 min gradient, including sample loading and column equilibration times. Data were acquired in data-independent mode using Xcalibur 4.5 software. MS1 Spectra were measured with a resolution of 120,000, an AGC target of $3 \times 10^6$ and a mass range from 350 to 1600 m/z. Per MS1, 29 equally distanced, sequential segments were triggered at a resolution of 30,000, an AGC target of $3 \times 10^6$, a segment width of 43 m/z, and a fixed first mass of 200 m/z. The stepped collision energies were set to 22.5, 25, and 27.

Two separate searches were conducted for the two digestion conditions. All DIA data were analyzed with Spectronaut software (*63*) (v18.6) using directDIA analysis methodology against a combined reference database including the *E. coli* proteome (NCBI RefSeq assembly GCF_000005845.2), T5 phage proteome (NCBI RefSeq assembly GCF_000858785.1), and the *Kpn*DRT2 RT and Neo (5 repeat) sequences. Cysteine carbamidomethylation was set as a fixed modification, and methionine oxidation and N-terminal acetylation were set as variable modifications. For the Neo-targeted experiment, trypsin, glu-c, and chymotrypsin were set as the digestion enzymes. For the global proteomics experiment, trypsin was set as the digestion enzyme. Normalization was performed using 'automatic normalization' in Spectronaut. Imputation was performed using 'global imputation' in Spectronaut for the global proteomics experiment, and was not performed for the Neo-targeted experiment. For differential protein abundance analysis, calculation of $\log_2$(fold change) and q-value was performed by Spectronaut using three independent biological replicates for each condition.

Infection time course:

For time course experiments assessing phage titer and concatenated RNA production during T5 infection of DRT2-expressing cells, *E. coli* MG1655 cells transformed with plasmids encoding WT or catalytically inactive (YCAA) *Kpn*DRT2 were grown to $OD_{600}$ of 0.4 in 25 ml volume. At the start of the experiment, 1 mL of culture was taken as the t = 0 (uninfected) time point for RNA extraction, and then phage lysate was added at MOI of 5. Cultures were incubated with shaking at 37 °C for 2 hours. Every 20 minutes, 200 µL and 1 mL volumes were taken for phage titer measurements and RNA extraction, respectively.

*Phage titer measurements*:

Phage titer measurements were taken over the course of T5 infection by removing 200 µL of culture from the ongoing infection and adding chloroform (5% final concentration) in order to completely lyse cells. Lysates were then centrifuged at 13,000 x *g* for 5 min in order to pellet cell debris. Enumeration of plaque forming units (PFU) was performed using the same plaque assay protocol as described above.

*RT-qPCR:*

Samples for RT-qPCR analysis were prepared with three independent biological replicates and were collected every 20 minutes for 2 hours after infection with T5 at MOI of 5, as described above. At each timepoint, 1 mL volumes of bacterial culture were removed and centrifuged at 3,000 x *g* for 3 min. The supernatant was removed, and the resulting pellet was resuspended in 750 µL of TRIzol and incubated at room temperature for 5 min. 150 µl of chloroform were added, and samples were mixed by shaking and centrifuged at 12,000 x *g* for 15 min at 4 °C. The upper aqueous phase was transferred to a new tube and mixed with an equal volume of absolute ethanol. Total RNA was purified using the Monarch RNA Cleanup Kit (NEB) and stored at -80 °C.

cDNA synthesis was performed using 500 ng of total RNA as the input, which was first treated with 1 µl of dsDNase (Thermo Fisher Scientific) in 1× dsDNase reaction buffer in a final volume of 10 µL, and incubated at 37 °C for 2 min. Reactions were stopped by adding DTT to a final concentration of 10 mM and heating to 55 °C for 5 min. Reverse transcription was performed using the iScript cDNA Synthesis Kit (BioRad) following the manufacturer's instructions. The samples were stored at -20 °C.

Quantitative PCR was performed in 10 µL reactions containing 5 µL SsoAdvanced Universal SYBR Green Supermix (BioRad), 0.5 µL of each primer pair at 10 µM concentration, and 4 µL of 25-fold diluted cDNA. Primers were designed to span the cDNA repeat junction. For normalization, primer pairs that anneal to the reference gene *rrsA* were used. Reactions were prepared in 384-well PCR plates (BioRad), and measurements were performed on a CFX384 RealTime PCR Detection System (BioRad) using the following thermal cycling parameters: polymerase activation and DNA denaturation (98 °C for 2.5 min), 40 cycles of amplification (98 °C for 10 s, 62 °C for 20 s), and terminal melt-curve analysis (decrease from 95 °C to 65 °C in 0.5 °C/5 s increments). Values are plotted as abundance of concatenated RNA, relative to *rrsA*, relative to the WT sample at t = 0 ($2^{-\Delta\Delta Cq}$). All primer sequences are provided in **Supplementary Table S4**.

*Northern blotting:*

RNA samples collected for RT-qPCR analysis, described above, were also used for Northern blotting analysis. After RNA purification by TRIzol and the Monarch RNA Cleanup Kit, samples were treated with TURBO DNase in TURBO DNase buffer (Thermo Fisher Scientific) for 30 min at 37 °C. Reactions were cleaned up using the Monarch RNA Cleanup Kit, and RNA concentrations were measured using the DeNovix RNA Assay.

Northern blotting was performed as previously described (*64*), with modifications. In brief, equal amounts of RNA (1.2 µg) were adjusted to 8 µL total volume with water and combined with 22 µL denaturing mix (15 µL formamide, 5.5 µL formaldehyde, and 1.5 µL 10× MOPS). Samples were heated at 55 °C for 15 min prior to separation on a denaturing agarose gel (1% agarose, 3.7% formaldehyde, 1× MOPS buffer) for 2.5 hours at 80 V. RNA was transferred to a Hybond-N+ membrane (GE Healthcare) by upward capillary transfer in 10× SSC (1.5 M NaCl, 0.15 M trisodium citrate dihydrate, pH 7). The next day, RNA was crosslinked to the membrane using a UV crosslinker, and the membrane was pre-hybridized in ULTRAhyb-Oligo buffer (Thermo Fisher Scientific) for 1 hour at 42 °C. A biotinylated oligonucleotide probe specific for the concatenated RNA repeat–repeat junction was added to the hybridization buffer at a final concentration of 5 nM and hybridization was performed overnight at 42 °C. The next day, the membrane was washed twice with Wash Buffer 1 (2× SSC with 0.1% SDS) and twice with Wash Buffer 2 (0.1× SSC with 0.1% SDS). The membrane was developed using the Chemiluminescent Nucleic Acid Detection Module Kit (Thermo Fisher Scientific) and imaged with an Amersham Imager 600 (GE Healthcare). The membrane was then stripped using boiling 0.1% SDS, pre-hybridized with ULTRAhyb-Oligo buffer, and reprobed using a biotinylated oligonucleotide probe specific for 16S rRNA. Hybridization, washes, and imaging were done as before. All probe sequences are provided in **Supplementary Table S4**.

Infection response growth curves:

Overnight cultures of *E. coli* MG1655 cells transformed with either empty vector (EV) or WT DRT2 expression vector were diluted 1:100 in LB with chloramphenicol (25 µg mL$^{-1}$), grown to exponential phase, and normalized to $OD_{600}$ of 0.2. 180 µL of cell culture were transferred into wells of a 96-well optical plate containing 20 µL of T5 lysate diluted to result in a final MOI of 5 or 0.05, or 20 µL of LB for the uninfected condition. The plate was incubated for 5 hours at 37 °C with shaking. $OD_{600}$ values were recorded every 10 minutes using a Synergy Neo2 microplate reader (Biotek).

Resazurin cell viability assays:

Cell viability was evaluated with the resazurin-based reagent alamarBlue HS (Thermo Fisher Scientific). 200 µL cultures were prepared as described above for cell growth experiments performed at varying MOIs. Infections proceeded for 3 hours before 180 µL of cell culture were mixed with 20 µL of alamarBlue HS and incubated with shaking at 37 °C. During incubation, fluorescence was measured in relative fluorescence units every 10 min according to the manufacturer's guidelines, using a Synergy Neo2 microplate reader (Biotek) with a monochromator module set to a fixed gain setting of 75. The fluorescence of blank LB controls was subtracted as background from all other measured values.

Neo induction experiments:

*Cellular growth curves*

*E. coli* MG1655 cells were transformed with plasmids encoding various repeat lengths of WT or mutant Neo. Individual colonies were inoculated in LB supplemented with kanamycin (50 µg mL$^{-1}$) and glucose (2%) and grown until cells reached $OD_{600}$ 0.8-1.0. The cells were then pelleted and resuspended in LB media with kanamycin (50 µg mL$^{-1}$). For each sample, $OD_{600}$ density was normalized to 0.1, and 200 µL of cell suspension were transferred to a 96-well clear-bottom plate. The $OD_{600}$ was measured using a Synergy Neo2 microplate reader (Biotek) while shaking at 37 °C for 50 min (until $OD_{600}$ reached ~0.3). Neo expression was then induced by the addition of arabinose (final concentration 0.5%) and theophylline (final concentration 0.5 mM), and cell growth was monitored for another 2 hours. For experiments testing the induction of diverse Neo homologs, growth rates were calculated using the formula The time window of 30 minutes to 80 minutes after induction was used to calculate the growth rate for each condition.

*Spot assays and CFU counting after Neo induction:*

To assess cell viability after *Kpn*Neo induction, a small volume from each well of the growth curve experiment described above was taken for plating on LB agar. 10× serial dilutions of each culture were prepared and spot-plated on LB agar supplemented with either kanamycin (50 µg $mL^{-1}$) and glucose (2%), or kanamycin (50 µg $mL^{-1}$), arabinose (0.5%), and theophylline (0.5 mM). Plates were incubated overnight at 37 °C, and colony forming units per milliliter (CFU/mL) were counted the next day.

Protein secondary structure prediction:

Six Neo protein sequences were aligned with MAFFT (*65*) (LINSI option; v7.520) , and the resulting alignment was visualized with Jalview (*66*) (v2.11.3.2). Secondary structural elements were predicted by submitting the alignment to Ali2D (*67*). The consensus predicted structure annotations and mean confidence values are plotted above the alignment in **Figure 5C**.

Protein tertiary structure prediction:

The Neo 3D structure was modeled using three independent prediction tools. The primary amino acid sequence of *Kpn*Neo was used as input for AlphaFold2 using MMseqs2 (via ColabFold), (*68*, *69*) and the same sequence was used as input for ESMFold (*70*). A multiple sequence alignment (MSA) of the Neo homologs shown in **Figure 5C** was used as input for trRosetta (*71*). All predictions were based on 3 concatenated repeats of Neo.

Start codon prediction:

The *Kpn neo* start codon was predicted using the RBS Calculator tool (*72*), using one cDNA repeat unit as the input sequence and specifying *K. pneumoniae* as the host organism.

Sequence identity matrices:

Pairwise sequence identity matrices were generated in Geneious from MAFFT alignments of ncRNA and cDNA nucleic acid sequences, or of RT and Neo amino acid sequences, using default settings. Accession numbers for RT proteins are listed in **Supplementary Table S5**.

RNA secondary structure prediction

*Kpn*DRT2 ncRNA secondary structure prediction (from single-sequence input) was performed using RNAfold (*24*) and visualized using RNAcanvas (*73*).

ncRNA covariance modeling:

Homologs of *Kpn*DRT2 were identified using the RT amino acid sequence (WP_012737279.1) as the seed query in a BLASTP search of the NR protein database (max target sequences = 100). Nucleotide sequences 1 kb upstream and downstream of *RT* genes were retrieved, clustered at 99.9% sequence identity to remove replicates using CD-HIT (*74*) (v4.8.1), and aligned using MAFFT (*75*) (v7.505). The resulting alignment was trimmed at the 5′ and 3′ ends to the exact boundaries of the ncRNA, as determined by RIP-seq experiments with *Kpn*DRT2. These putative ncRNA sequences were clustered at 95%

sequence identity using CD-HIT and realigned using mLocARNA (*76*) (v1.9.1) with default parameters. The resulting structure-based multiple sequence alignment was used to build and calibrate a covariance model (CM) using the Infernal suite (*77*) (v1.1.4). The CMsearch function of Infernal was then used to scan through nucleotide sequences of additional *drt2* loci and 1-kb flanking regions, generated by an expanded BLASTP search (max target sequences = 5000) queried on *Kpn*DRT2 and clustered at 85% sequence identity using CD-HIT. The final hits (n = 303 DRT2 loci, including *Kpn*DRT2) from the CM used to identify *Kpn*DRT2-like ncRNAs were evaluated for statistically significant co-varying base pairs with R-scape (*78*) at an *E*-value threshold of 0.05 (**fig. S2D**).

Phylogenetic analyses:

An initial set of DRT2 sequences was identified by querying the *Kpn*RT protein sequence (WP_012737279.1) against the NR database with PSI-BLAST (3 iterations; default settings) (*79*). The top 500 results from this search did not produce any clusters at a threshold of 80% amino acid identity, so this diverse set of homologs was used for an additional BLASTP (-evalue 0.01 -max_target_seqs 1000000) search of a local copy of the NCBI NR database (downloaded on April 4, 2023). The resulting hits were further restricted to an e-value cutoff of $1 \times 10^{-30}$, resulting in a set of 3,056 protein accessions, for which identical protein group (IPG) information was pulled from NCBI with the Batch Entrez tool. Where possible, two genomes encoding each unique DRT2 homolog were randomly sampled from the IPG information, and these genomic sequences were retrieved from NCBI with the Batch Entrez. DRT2 homologs for which we were unable to retrieve IPG information or genomic sequences were removed from the analysis, resulting in a final dataset of 2,116 DRT2 homologs (**Supplementary Table S1**). 616 protein sequences in this final DRT2 dataset were also identified as DRT2 homologs in a previous analysis of reverse transcriptases (*19*), while no other non-DRT2 homologs from that previous analysis were present in our dataset. Finally, this set of DRT2 sequences was aligned with MAFFT (LINSI option; v7.520) and a phylogenetic tree was constructed with FastTree (*80*) (-wag -gamma; 2.1.11), before being visualized with iTOL (*81*). A subtree of KpnDRT2-like sequences was constructed by manually subsetting the tree in **fig. S6A** to include only a monophyletic clade encompassing KpnDRT2. Eight sequences with unexpectedly long branch lengths were manually pruned from this subtree, resulting in a phylogeny of 539 KpnDRT2-like sequences (**Fig. 5B**).

Systematic ncRNA and Neo prediction:

An updated *Kpn*DRT2-like CM (v2) was built by retrieving genomes for the top 500 DRT2 hits from the PSI-BLAST search described above, extracting the DRT2 loci (*drt2* +/- 1 kb), searching each locus with the CMsearch function of Infernal (v1.1.5; default parameters), aligning hits that met the inclusion threshold (n = 287) with LocARNA (v2.0.0; mlocarna option with default parameters), and building a CM with the CMbuild function of Infernal (v1.1.5; default parameters).

DRT2 loci, corresponding to the *RT* gene and 1 kb of upstream and downstream sequence, were then extracted from genomes encoding the 2,116 DRT2 homologs described above, using coordinates in the IPG dataset. To identify putative ncRNA sequences, these loci were queried with the *Kpn*DRT2-like ncRNA CM (v2) using the CMsearch function of Infernal (v1.1.5), with default parameters. Hits that met the inclusion threshold (e-value < 0.01) were extracted using the coordinates in the CMsearch output, and these putative ncRNA sequences were de-duplicated, prior to alignment of the sequences with the DECIPHER package (*82*) (v2.30.0) in R. The *Kpn*DRT2 locus was used as a reference to extract likely cDNA regions from the resulting alignment.

To predict Neo sequences in homologous DRT2 loci, the reverse complements of putative cDNA sequences were assessed in all three possible reading frames to determine which frame contained the fewest stop codons; these were assumed to be the *neo* open reading frames (ORFs). Start codons (ATG, GTG, TTG) were then probed in the resulting putative *neo* ORFs, and the start codon of *neo* was presumed to occur after the first ten amino acids of the putative reading frame translation, consistent with the *Kpn*DRT2 locus. Putative Neo sequences were then constructed by concatenating the translation of this downstream start codon in the putative *neo* ORF through the end of the putative cDNA (which represents the first unit of cDNA produced by the putative DRT2 rolling circle mechanism; repeat 1), to a translation of the full length putative *neo* ORF (which represents cDNA units produced by successive rounds of the DRT2 rolling circle mechanism; repeat 1 + *n*). Putative *neo* sequences that did not contain an internal stop codon, were then checked to determine if the final ten amino acids of the Neo sequence (i.e., translated from repeat 1+*n*) were identical to the final ten amino acids of Neo translated from the first unit of cDNA synthesis (i.e., translated from repeat 1) (**Fig. 5A**). Putative Neo sequences that met this criterion were predicted to represent bona fide Neo protein products of DRT2 immune systems.

Putative ncRNA sequences identified with the *Kpn*DRT2-like ncRNA CM (v2) were primarily restricted to a monophyletic clade that included *Kpn*DRT2 (**Fig. 5B**). An alignment of these ncRNA sequences was built with the DECIPHER package in R, and sequence logos (**fig. S6B**) were generated with the web-based version of WebLogo (***83***). Logos of cDNA sequences with identified Neo proteins (**fig. S6B**) were similarly built from an MSA generated with DECIPHER. To identify ncRNAs in other regions of the larger phylogenetic tree presented in **fig. S6A**, additional CMs were constructed via the same approach described above (i.e., mlocarna with default settings, CMbuild in Infernal) by manually selecting regions of the tree that had CM hits for at least three closely related DRT2 sequences. These CMs were used to search the DRT2 loci, and then new CMs were built from the resulting hits; this process was iterated until ncRNAs had been identified across most DRT2 systems. Exemplary ncRNA CMs generated in this process are shown in **fig. S6C**. Finally, Neo sequences were predicted in these newly identified putative ncRNAs using the same approach described above.

### SUPPLEMENTARY FIGURES

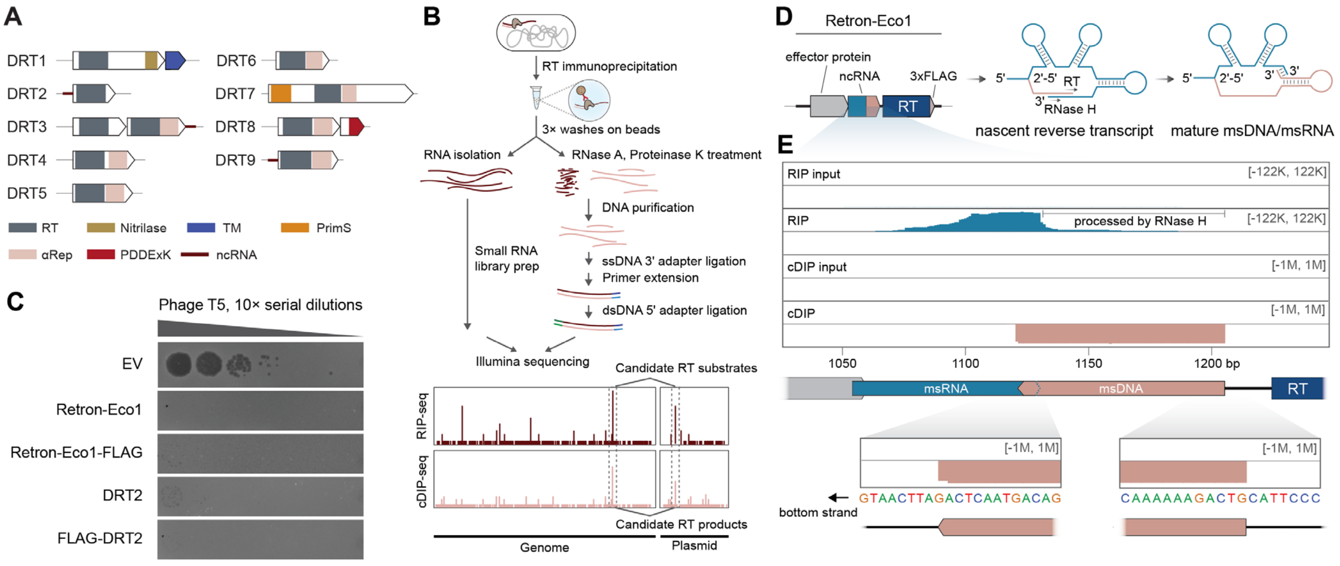


**Fig. S1. Overview and validation of cDNA immunoprecipitation and sequencing (cDIP-seq) approach using Retron-Eco1. (A)** Summary of operonic configurations for DRT systems with experimentally validated phage defense activity. Protein domains and associated ncRNAs are indicated. **(B)** Detailed overview of combined RIP-seq and cDIP-seq workflow (Materials and Methods). **(C)** Plaque assay showing that FLAG-tagged RT proteins in Retron-Eco1 (formerly Ec86) and *Kpn*DRT2 systems retain WT defense activity. **(D)** Schematic of the Retron-Eco1 operon (left), retron-encoded ncRNA (middle), and the nascent and mature multi-copy single-stranded DNA (msDNA) products resulting from reverse transcription and RNase H processing steps (right). **(E)** RIP-seq and cDIP-seq coverage tracks for Retron-Eco1, plotted alongside corresponding input controls over the plasmid-encoded Retron-Eco1 locus. The drop-off in RIP coverage is consistent with processing of the msRNA by RNase H. The magnified insets highlight the accuracy and single-nucleotide precision of cDIP-seq in identifying the msDNA, which are consistent with previous reports (*21*).

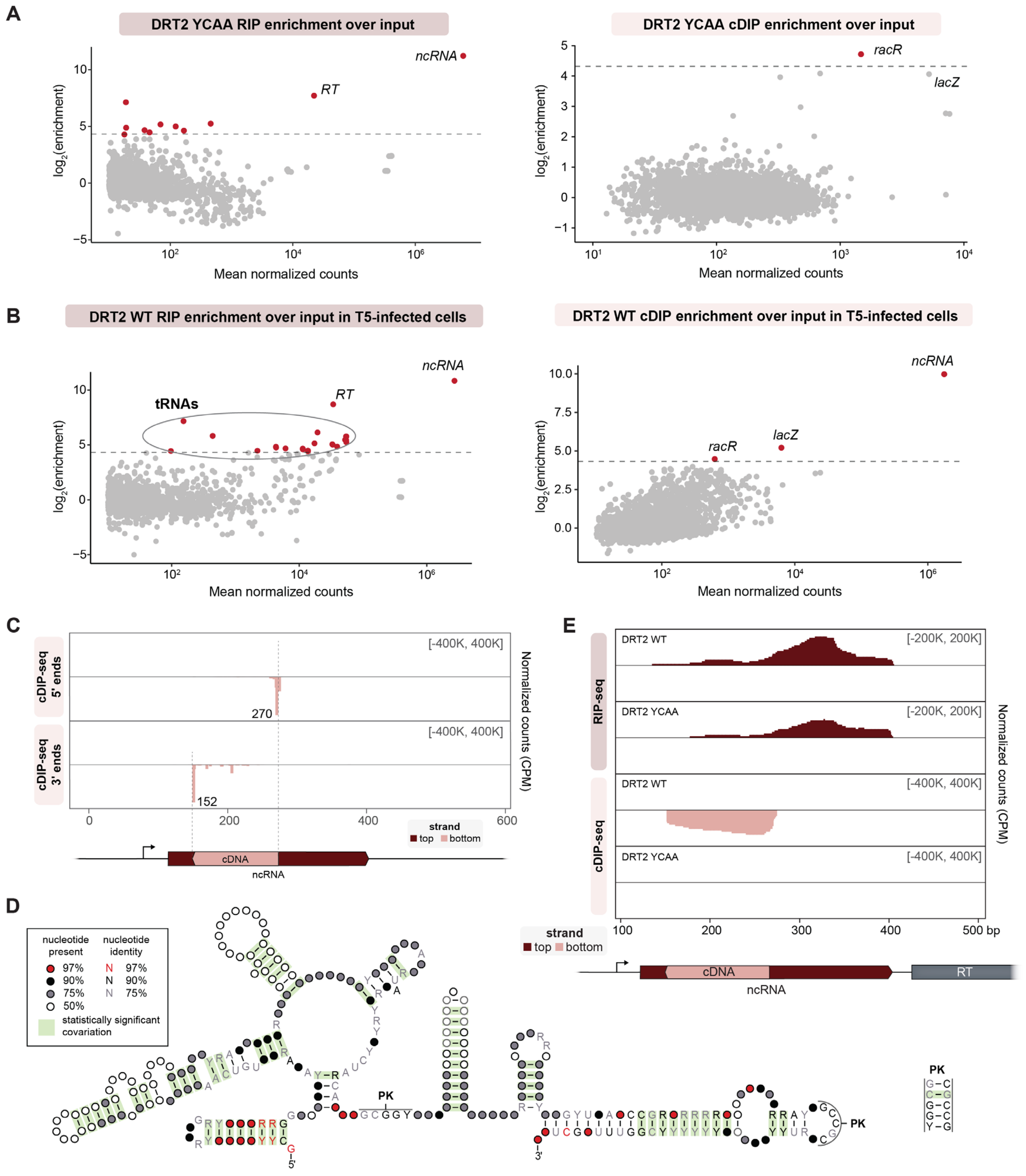


**Fig. S2. Additional analyses of DRT2 RIP-seq and cDIP-seq data. (A)** MA plots showing catalytically inactive RT (YCAA)-mediated enrichment of RNA (left) and DNA (right) loci from RIP-seq and cDIP-seq experiments, relative to input controls. Each dot represents a transcript, and red dots denote transcripts with > 20-fold enrichment and false discovery rate (FDR) < 0.05. The cDIP-seq enrichment of *lacZ* and *racR* with an RT-inactive mutant indicates that they do not represent true cDNA synthesis products. **(B)** MA plots as in **A** for RIP-seq and cDIP-seq of WT DRT2 from T5 phage-infected cells. Transcripts from the *ncRNA* locus are similarly enriched as in RIP-seq and cDIP-seq experiments from uninfected cells (**Fig. 1B**), but we also observed enrichment of diverse tRNAs in the RIP-seq dataset. **(C)** Histogram of mapping coordinates for 5′ and 3′ ends of cDIP-seq fragments visualized over the DRT2 ncRNA locus (bottom). Coordinates are numbered from the beginning of the *K. pneumoniae*-derived sequence on the DRT2 expression plasmid; the precise coordinates are indicated next to their corresponding peaks. (**D**) Covariance model for ncRNA sequences from *Kpn*DRT2 and related loci (n = 303). PK denotes a predicted pseudoknot interaction between the indicated regions. **(E)** RIP-seq and cDIP-seq coverage tracks for either WT RT or a catalytically inactive RT mutant (YCAA) from T5 phage-infected cells. Red and pink denote top and bottom strands, respectively, and the DRT2 locus is shown at bottom. Data are normalized for sequencing depth and plotted as counts per million reads (CPM); coordinates are numbered as in **C**.

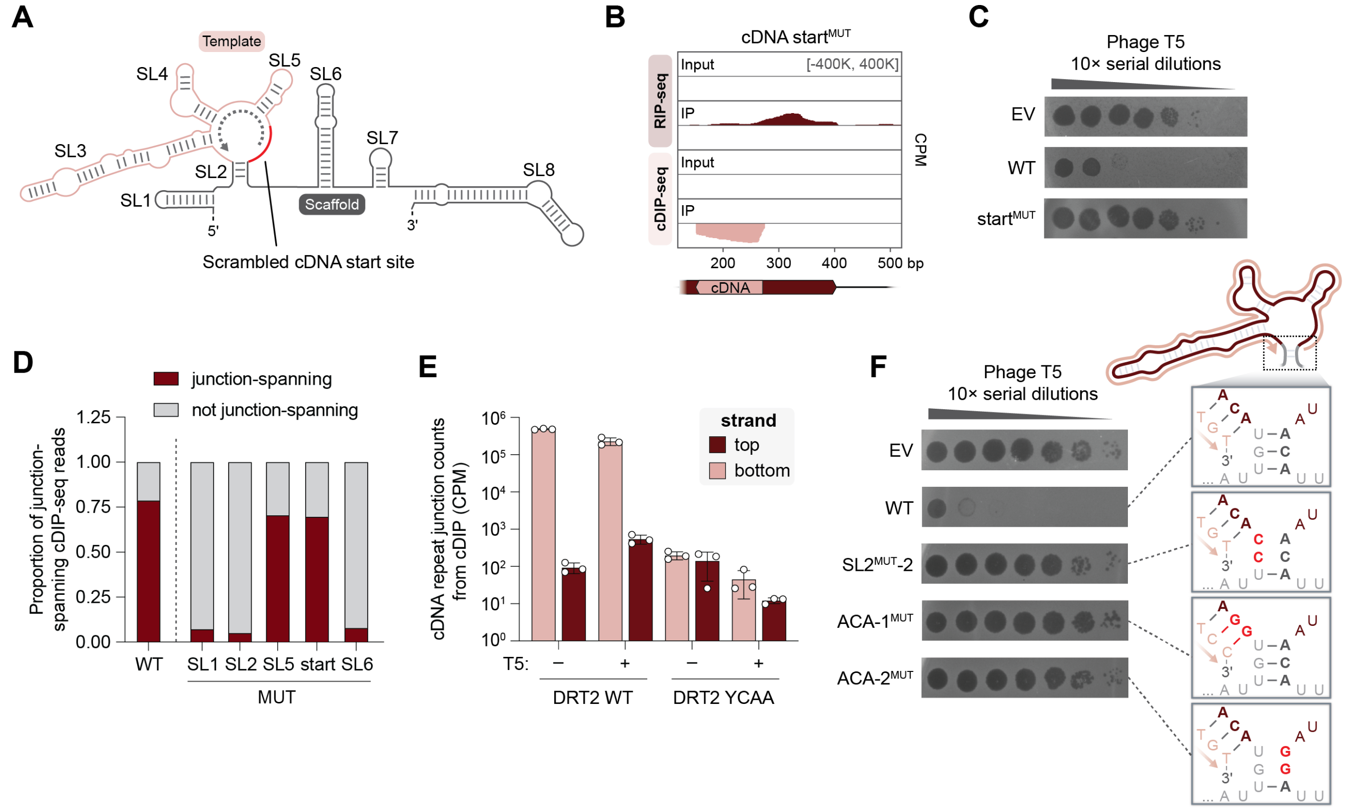


**Fig. S3. Molecular dissection of phage and ncRNA requirements during rolling-circle reverse transcription (RCRT). (A)** Schematic of the DRT2 ncRNA secondary structure, indicating the region surrounding the reverse transcription start site (red line) that was mutated for RIP-seq, cDIP-seq, and phage defense experiments. **(B)** RIP-seq and cDIP-seq coverage tracks for the scrambled cDNA start site mutant (cDNA start$^{MUT}$) alongside input controls, showing RNA binding and cDNA synthesis activities comparable to WT (**Fig. 1C**). **(C)** Plaque assay demonstrating loss of phage defense activity with the cDNA start site mutant. **(D)** Stacked bar graph quantifying cDIP-seq reads mapping across the cDNA repeat–repeat junction as a proportion of total reads mapping to the DRT2 ncRNA locus, for the WT system and indicated ncRNA SL mutants in uninfected cells. The data demonstrate that programmed template jumping requires an intact SL2 alongside conserved ncRNA features at the 5′ end (SL1) and within the scaffold region (SL6). **(E)** Bar graph quantifying the abundance of junction-spanning reads from cDIP-seq experiments with the indicated conditions. Red and pink denote top and bottom strands, respectively; data are mean ± s.d. (n = 3). Based on the difference between these results and those obtained from total DNA sequencing (**Fig. 2E**), we conclude that the RT likely releases double-stranded cDNA after second-strand synthesis, leading to a lack of enrichment for the top strand in these experiments. **(F)** Plaque assay showing loss of phage defense activity with an additional SL2 mutant (SL2$^{MUT}$-2), as well as with mutations to either ACA-1 or ACA-2 (see **Fig. 2G**). Schematics of mutants are shown in the insets at right. Template nucleotides are in maroon, cDNA nucleotides are in pink, and ACA motifs are bolded. Mutated nucleotides are bolded in bright red.

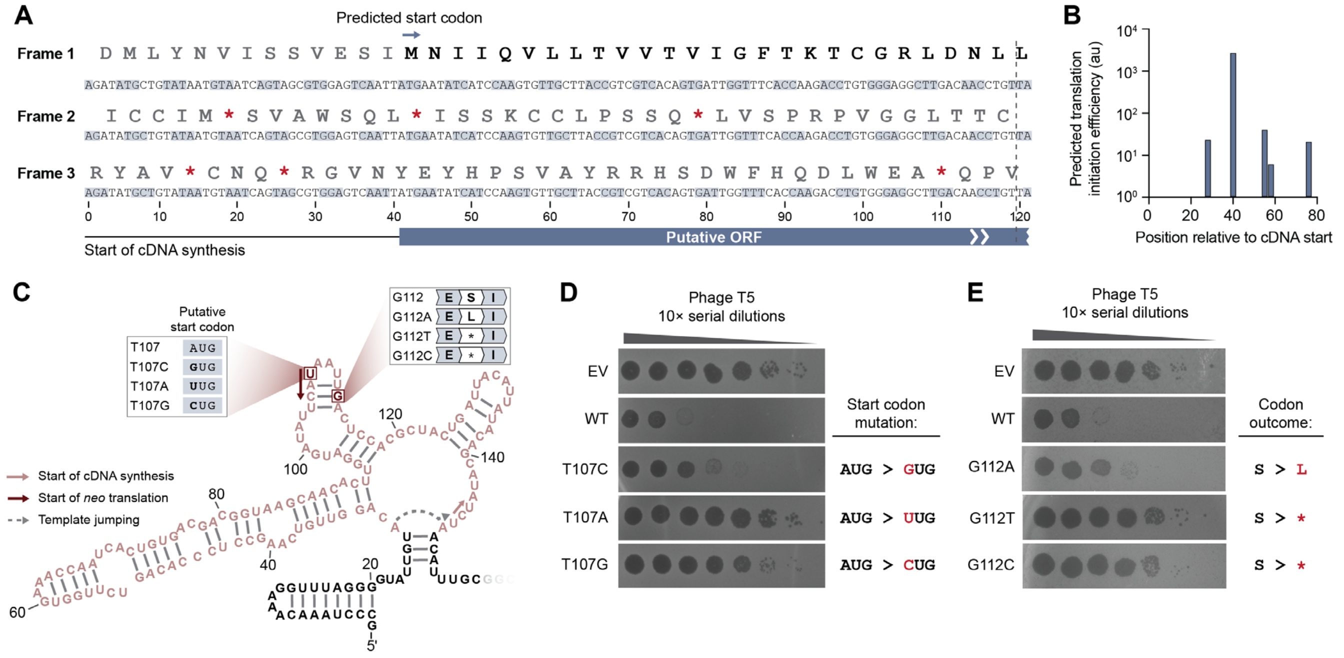


**Fig. S4. Additional genetic evidence that the DRT2 concatenated cDNA encodes a functional open reading frame (ORF).** **(A)** *In silico* translation of the cDNA repeat sequence in all three possible reading frames, demonstrating that Frame 1 lacks stop codons. **(B)** Prediction of translation initiation efficiency for an mRNA derived from one cDNA repeat, based on an *in silico* ribosome binding site (RBS) calculation tool (*72*). Predicted values are shown in arbitrary units (au) for each potential AUG start codon, as well as the non-canonical start codons GUG and UUG, found within the mRNA. **(C)** Schematic of the cDNA template region (pink), with experimentally tested mutations indicated. Single-bp substitutions were designed to either disrupt the putative start codon or introduce a missense or stop codon after one near-full-length *neo* repeat (39/40 amino acids). **(D)** Plaque assay demonstrating that mutation of the putative AUG start codon to GUG, a common non-canonical start codon in *E. coli* (*84*), is partially tolerated, whereas mutations to UUG and CUG result in a complete loss of defense. **(E)** Plaque assay demonstrating that a single-bp substitution at G112 is partially tolerated when resulting in a missense codon, but abolishes defense activity when resulting in a stop codon.

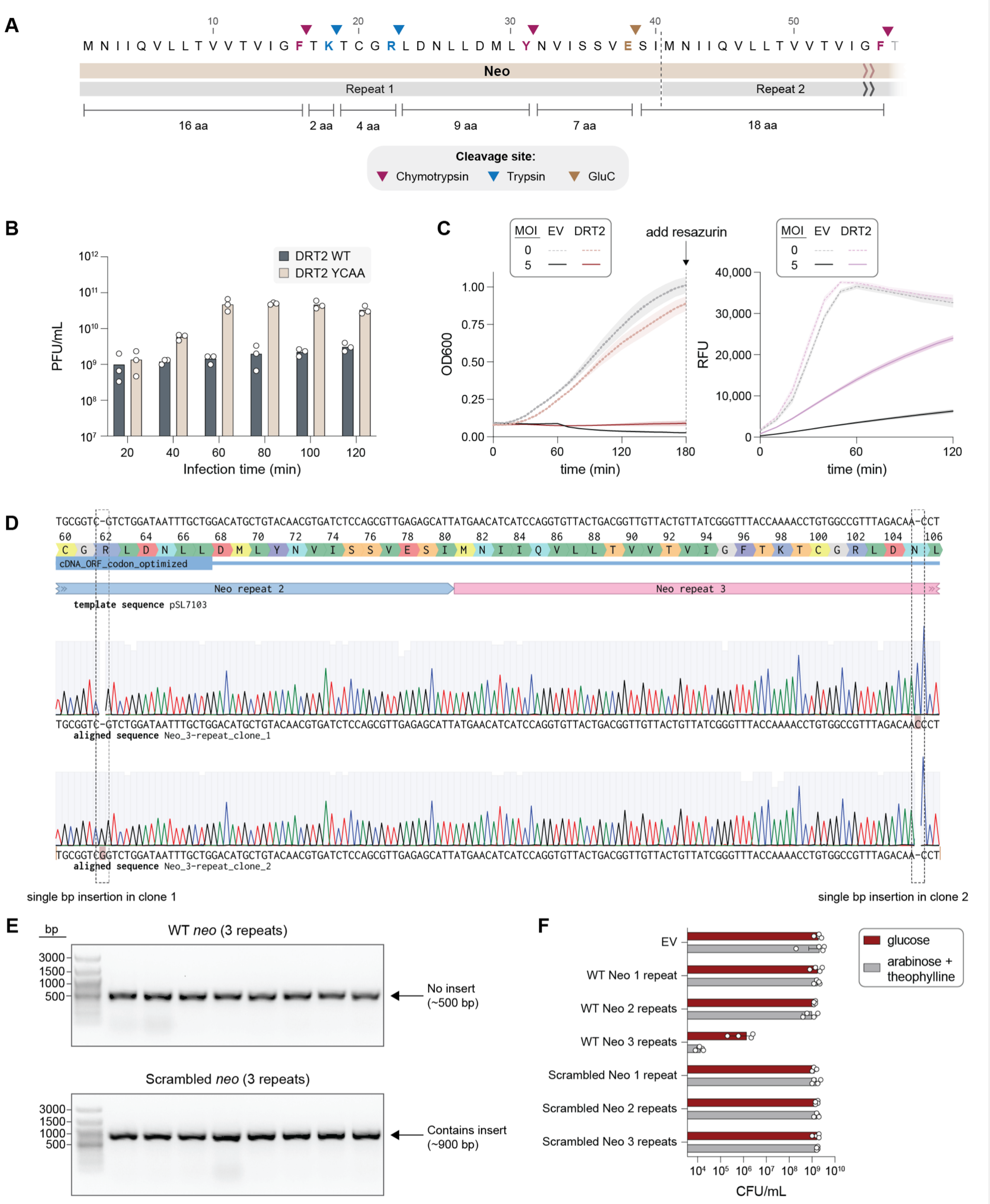


**Fig. S5. Detection and recombinant expression of *neo*. (A)** Cleavage map for Neo digestion using a custom 3-enzyme protease cocktail prior to liquid chromatography with tandem mass spectrometry (LC-MS/MS) analysis. A full Neo repeat is shown, together with the portion of the downstream repeat leading up to the next cleavage site. Cleavage sites are indicated with triangles, colored by enzyme; the ideal peptide fragment size for MS is 9–15 amino acids. **(B)** T5 phage titer measurements at the indicated time points after a high-MOI infection of cells expressing DRT2 (WT or YCAA). Data are shown as mean ± s.d. (n = 3). **(C)** *Left*: Growth curves of strains transformed with empty vector (EV) or the WT DRT2 system, +/- T5 phage infection at an MOI of 5. *Right*: Relative fluorescence unit (RFU) measurements after addition of resazurin to the same cultures after 3 hours of growth. The shaded regions indicate the standard deviation across independent biological replicates (n = 6). **(D)** Sanger sequencing traces showing the presence of frameshift mutations in *neo* when attempting to isolate clones of a 3-repeat *neo* expression vector. **(E)** Colony PCR demonstrating unsuccessful cloning of 3-repeat WT *neo* into a standard expression vector, as compared to successful cloning of 3-repeat scrambled *neo* mutant. For each experiment, 8 clones were randomly selected and subjected to PCR, which is expected to generate a ~900-bp band with the insert. **(F)** Colony forming unit (CFU) measurements for cultures from **Fig. 4H** plated on repressor (2% glucose) or inducer (0.5% arabinose and 0.5 mM theophylline) after the final $OD_{600}$ measurement time point.

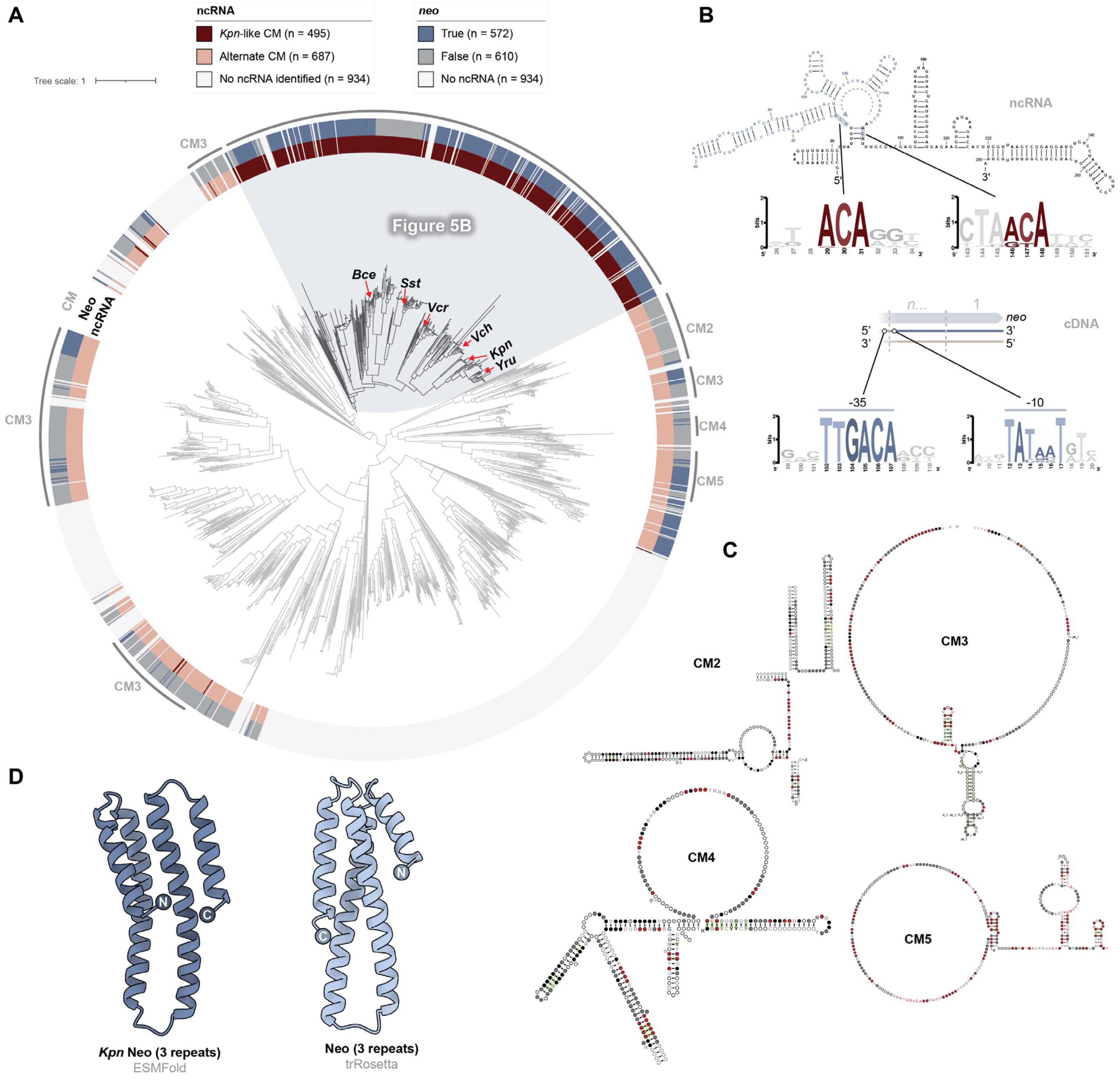


**Fig. S6. Widespread presence of *neo* across diverse clades of DRT2 systems. (A)** Phylogenetic tree of DRT2-encoded RT homologs (n = 2,116). The rings depict ncRNA sequences identified via CM searches (inner) and the presence of bioinformatically identified *neo* genes (outer). The subtree shown in **Figure 5B** comprises the sequences highlighted in light blue. **(B)** *Top*: Conservation of ACA-1 and ACA-2 motifs that mediate programmed template jumping (n = 257 loci). *Bottom*: Conservation of -35 and -10 promoter elements flanking the *neo* repeat junction (n = 203 loci). **(C)** Exemplary covariance models for the DRT2 ncRNA derived from additional clades in **A**; the locations of each CM within the phylogenetic tree are indicated. **(D)** Additional *in silico* predictions of the 3D structure of a 3-repeat Neo polypeptide. *Kpn* Neo was used as a single-sequence input for ESMFold (*70*), while an MSA of the homologs shown in **Fig. 5C** was used as input for trRosetta (*71*).

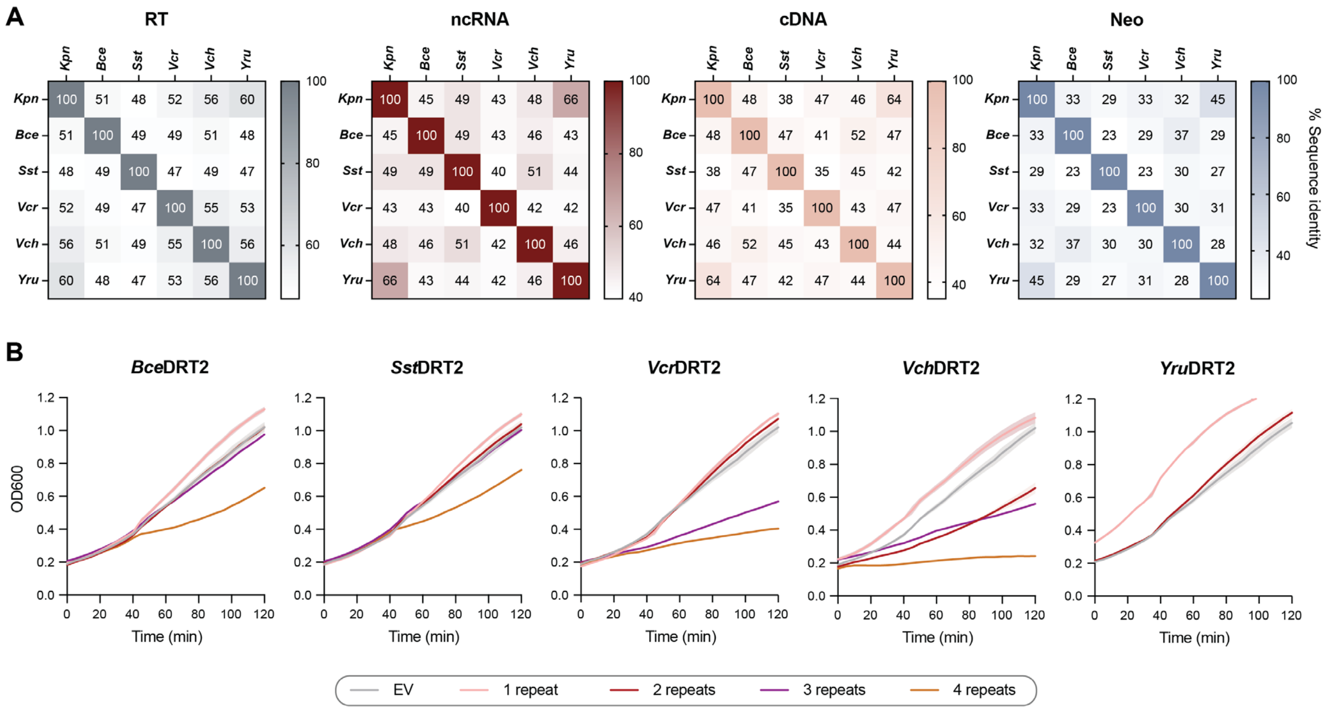


**Fig. S7**. **Diverse *neo* homologs induce repeat length-dependent growth arrest. (A)** Sequence identity matrices of the RT, ncRNA, cDNA repeat, and Neo polypeptide sequences for experimentally tested DRT systems. **(B)** Growth curves of strains transformed with *neo* homologs of the indicated repeat lengths, alongside an empty vector (EV) control (related to **Fig. 5G**). Expression was induced with 0.5% arabinose and 0.5 mM theophylline. Shaded regions indicate the standard deviation across independent biological replicates (n = 3). Note that cloning of *neo* from *Y. ruckeri* with 3 or 4 repeats was unsuccessful, presumably due to toxicity from even minimal leaky expression of the Neo protein.
